# *In vivo* single-cell CRISPR uncovers distinct TNF-α programs in clonal expansion and tumorigenesis

**DOI:** 10.1101/2023.07.13.548697

**Authors:** Peter F. Renz, Umesh Ghoshdastider, Simona Baghai Sain, Fabiola Valdivia-Francia, Ameya Khandekar, Mark Ormiston, Martino Bernasconi, Jonas A. Kretz, Minkyoung Lee, Katie Hyams, Merima Forny, Marcel Pohly, Xenia Ficht, Stephanie J. Ellis, Andreas E. Moor, Ataman Sendoel

**Affiliations:** Institute for Regenerative Medicine (IREM), University of Zurich, Wagistrasse 12, CH-8952 Schlieren-Zurich, Switzerland; Department of Biosystems Science and Engineering, ETH Zurich, Mattenstrasse 26, 4058, Basel, Switzerland; Life Science Zurich Graduate School, Molecular Life Science Program, University of Zurich/ ETH Zurich, Switzerland; Max Perutz Labs, Vienna BioCenter Campus (VBC), Dr.-Bohr-Gasse 9, 1030, Vienna AT; University of Vienna, Center for Molecular Biology, Department of Microbiology, Immunobiology, and Genetics, Dr.-Bohr-Gasse 9, 1030, Vienna, Austria

## Abstract

The tumor evolution model posits that malignant transformation is preceded by randomly distributed driver mutations in cancer genes, which cause clonal expansions in phenotypically normal tissues. Although clonal expansions occur frequently in human epithelia and can remodel almost entire tissues, the mechanisms behind why only a small number of clones transform into malignant tumors remain enigmatic. Here, we develop an in vivo single-cell CRISPR strategy to systematically investigate tissue-wide clonal dynamics of the 150 most frequently mutated squamous cell carcinoma genes. We couple ultrasound-guided in utero lentiviral microinjections, single-cell RNA sequencing, guide capture and spatial transcriptomics to longitudinally monitor cell type-specific clonal expansions, document their underlying gene programs and contrast clonal expansions from tumor initiation. We uncover a TNF-α signaling module that acts as a generalizable driver of clonal expansions in epithelial tissues. Conversely, during tumorigenesis, the TNF-α signaling module is downregulated, and instead, we identify a subpopulation of invasive cancer cells that switch to an autocrine TNF-α gene program. By analyzing clonally expanded perturbations and their frequency in tumors, we demonstrate that the autocrine TNF-α gene program is associated with epithelial-mesenchymal transition (EMT) and is preexistent in a subpopulation of expanded epidermal stem cells, contributing to the predisposition for tumor initiation. Finally, we provide in vivo evidence that the epithelial TNF-α gene program is sufficient to mediate invasive properties of epidermal stem cells and show that the TNF-α signature correlates with shorter overall survival in human squamous cell carcinoma patients. Collectively, our study demonstrates the power of applying in vivo single-cell CRISPR screening to mammalian tissues and unveils distinct TNF-α programs in tumor evolution. Understanding the biology of clonal expansions in phenotypically normal epithelia and the mechanisms governing their transformation will guide the development of novel strategies for early cancer detection and therapy.

## Introduction

Tumor evolution begins with the transformation of a single cell that expands to form a tumor mass. Central to the tumor evolution model is that transformation is preceded by randomly distributed driver mutations in cancer genes, which cause clonal expansions in phenotypically normal tissues. These clonal expansions increase the pool of cells that can accumulate additional driver mutations until a sufficient set of mutations and gene expression changes are acquired to enable malignant transformation. Notably, driver mutations in cancer genes occur with a surprisingly high frequency in normal human epithelia such as the skin, esophagus, bladder or colon^1^. For instance, *NOTCH1* mutations are prevalent in multiple epithelial tissues, including the esophagus, where their frequency even exceeds that observed in cancer^1,2^. Clonal expansion size does also not necessarily correlate with the capacity to induce malignant transformation, as illustrated by the presence of large clones with activating *FGFR3* mutations in human skin that rarely progress to malignancy^3^. In addition, rather than only being viewed as the underlying basis for transformation, highly competitive driver mutation clones in the esophagus can also play a tumor-suppressive role by eliminating emerging new tumors through clonal competition^4^. These observations raise the question of the fundamental relationship between clonal expansions in normal epithelia and their role in driving malignant transformation, leaving us with major gaps in our understanding of the earliest stages of tumorigenesis. Critical questions that remain unanswered include the gene expression programs and cell types underlying clonal expansion in normal epithelia and the molecular processes that distinguish them from malignant transformation. Answering these questions at a tissue-wide level *in vivo* remains a major challenge. Although pooled CRISPR screens are a powerful tool for studying clonal growth and inferring gene functions, they are limited to simple readouts such as proliferation, averaged over all tissue cell types. The systematic monitoring of genetically perturbed clones in normal epithelia and early stages of tumorigenesis requires scalable, tissue-wide platforms that can characterize the consequences of gene perturbations across all cell types.

Here, we use the mouse skin as a model to systematically monitor clonal expansion in epithelia and its role in tumor initiation. We combine ultrasound-guided *in utero* microinjections into embryonic stage (E) 9.5 embryos with an adapted CRISPR droplet sequencing (CROP-seq) strategy^5^ and spatial transcriptomics to longitudinally document the clonal expansion of the 150 most frequently mutated cancer genes at single-cell transcriptomic resolution. Our findings unravel the biology of clonal expansions in phenotypically normal epithelia and uncover distinct TNF-α gene programs in clonal expansion and tumor initiation.

## Results

### *In vivo* single-cell CRISPR screen to monitor clonal expansions of cancer genes

To monitor tissue-wide clonal expansion by an *in vivo* single-cell CRISPR strategy, we adjusted the previously reported CROP-seq system^5^. Unlike other single-cell CRISPR systems that rely on barcodes, the CROP-seq vector expresses the single-guide RNA (sgRNA) within a polyadenylated transcript, which circumvents concerns associated with barcode swapping due to lentiviral template switching^6^. To adapt the CROP-seq plasmid to our *in vivo* experimental requirements, we replaced the Puromycin resistance cassette in the original plasmid with mCherry to allow the selection of infected cells by fluorescence-activated cell sorting (FACS) (Figure 1a).

**Figure 1.**
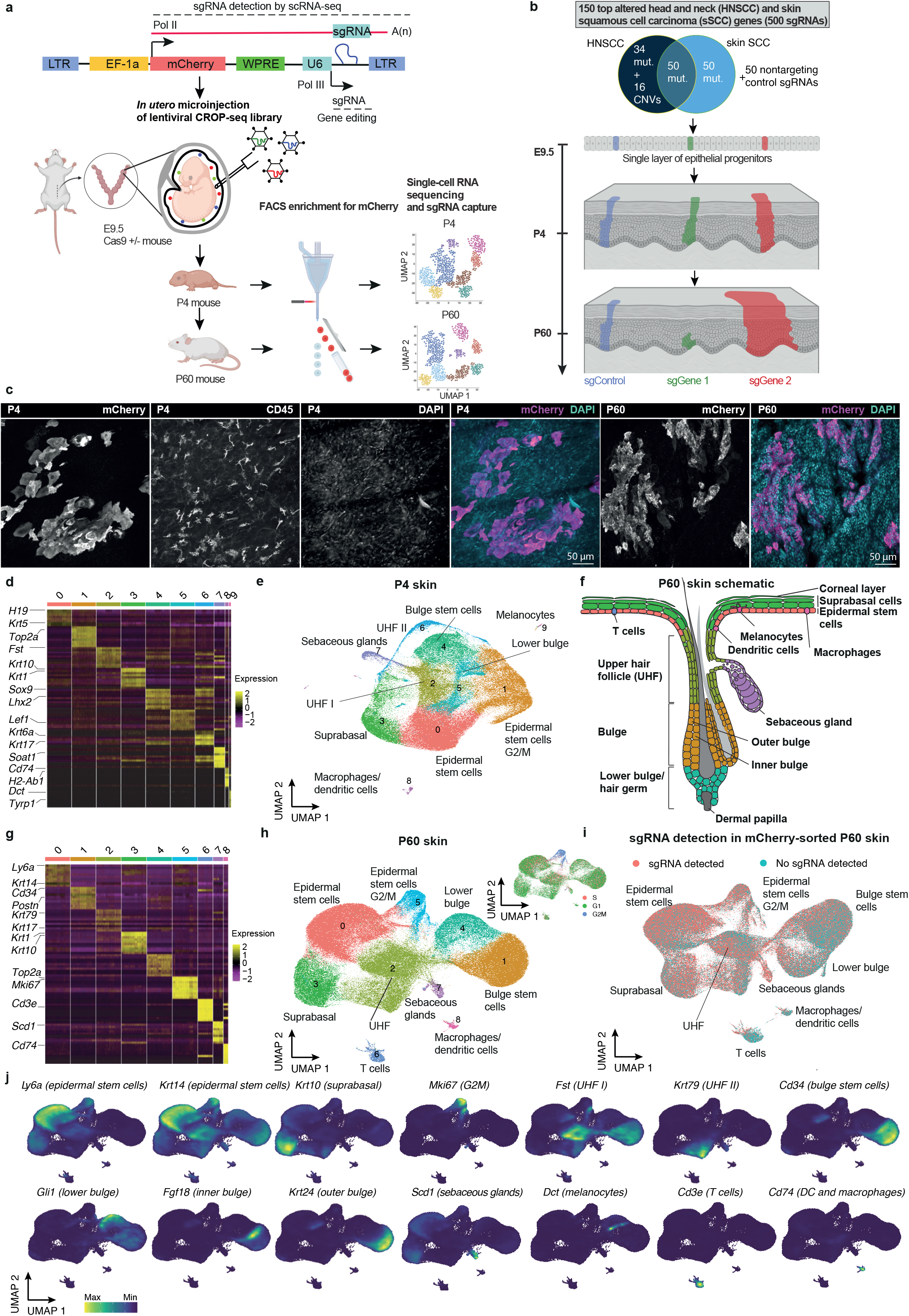
*In vivo* CRISPR screening to monitor clonal expansion in the mouse skin induced by mutations in cancer genes. **a**, Schematic outline of the *in vivo* CROP-seq strategy to couple pooled CRISPR screening with single-cell transcriptomic readout. A high-titer lentiviral library is introduced into *Rosa26-Cas9* E9.5 embryos via ultrasound-guided *in utero* microinjections, infecting a single layer of epidermal progenitors and stably propagating during development. Mouse epidermis from postnatal day 4 (P4) and 60 (P60) is harnessed, followed by FACS enrichment and subsequent single-cell RNA sequencing (scRNA-seq) with sgRNA capture. **b**, The sgRNA library consists of the 150 most frequently mutated genes in head and neck (HNSCC) and skin squamous cell carcinoma patients (sSCC). Three sgRNAs per gene were selected, along with 50 non-targeting control sgRNAs, resulting in a library of 500 sgRNAs. **c**, Whole-mount immunofluorescence staining of mCherry-positive clonal expansions in P4 and P60 mouse skin. Immunofluorescence images provide a planar view across the epidermis. CD45 staining marks hematopoietic cells. Scale bars, 50 **μ**m. **d**, Heatmap displaying the expression profiles of the top marker genes across individual mCherry-positive cells in P4 animals. **e**, Uniform manifold approximation and projection (UMAP) of the ten major cell populations identified by the *in vivo* CROP-seq strategy of P4 animals (n = 120,077 cells, from 58 animals, following filtering and sgRNA annotation). UHF, upper hair follicle. **f**, Schematic representation of the anatomical localization of the different cell populations in the P60 mouse skin. **g**, Heatmap displaying the expression profiles of the top marker genes across individual mCherry-positive cells in P60 animals. **h**, UMAP of the nine major cell populations identified by the *in vivo* CROP-seq strategy of P60 animals (n = 183,084 cells, from 10 animals, following filtering and sgRNA annotation). Inset depicts the cell cycle phase of the different cell populations. UHF, upper hair follicle. **i**, sgRNAs are uniformly detected in all major cell populations. The UMAP shows total mCherry-sorted P60 cells, divided into cells with detected sgRNA and cells without detected sgRNA. sgRNAs are captured by a dial-out PCR and sequenced together with the single-cell RNA sequencing library. **j**, Visualization of cell-type specific marker gene expressions by Nebulosa density plots, highlighting their specific localization in the P60 UMAP. DC, dendritic cell (Langerhans cell).

To longitudinally monitor the function of cancer genes, we chose the 150 most frequently altered genes in human head and neck (HNSCC) and skin squamous cell carcinomas (sSCC), comprising 134 mutated genes (predominantly tumor suppressors) and 16 copy number variations (CNVs) (Figure 1a-b, Extended Data Figure 1a-c). We selected 3 sgRNAs per gene and included 50 non-targeting control sgRNAs, resulting in a CROP-seq library of 500 sgRNAs used for high-titer lentivirus production. Leveraging the ultrasound-guided *in utero* microinjection system^7,8^, we delivered the lentiviral library into E9.5 embryos to infect the single-layered surface ectoderm, resident immune cells and melanoblasts of constitutively *Cas9*-expressing mouse embryos and introduce loss-of-function mutations in each of the selected cancer genes^9^. The library carrying the 500 sgRNAs is stably propagated by the surface ectoderm, which gives rise to several structures, including the skin epidermis, hair follicles, sebaceous glands and oral mucosa^7,8,10^. At postnatal day 4 (P4) and 60 (P60), we collected epidermal tissues, sorted for mCherry-positive infected cells, and subjected them to single-cell RNA sequencing (scRNA-seq) to assess gene expression profiles and read out the corresponding sgRNA identity (Figure 1a). We generated scRNA-seq libraries comprising a total of 477,773 single cells. Following stringent filtering and sgRNA annotation, we obtained 120,077 P4 and 183,084 P60 single cells, providing an average coverage of 240x P4 and 366x P60 cells per guide. 499 sgRNAs were detected throughout the T0 (representation in the initial library), P4 and P60 time points (p63 sgRNA3 was not detected at P4 and P60 due to a complete depletion). To reduce double sgRNA infections, we aimed at a multiplicity of infection (MOI) < 1, resulting in infection rates between 1-12.5 % in P4 animals (median 2.4%) and multiple infection rates under 1%^7^ (Extended Data Figure 1d). These low MOIs minimized the possibility of perturbed cells being influenced by neighboring cells with different perturbations. Immunofluorescence wholemount imaging of the P4 and P60 skin confirmed the clearly separated mCherry-positive clusters within the epidermis (Figure 1c).

To validate our approach, we first confirmed the efficiency of select sgRNAs and found a 67-86% editing efficiency in keratinocytes (Extended Data Figure 1e). Second, the control library of 50 sgRNAs was overall stably propagated and evenly distributed (Extended Data Figure 1f). Third, to validate the accuracy of our sgRNA assignments and single-cell transcriptomic data, we examined cells carrying sgRNAs targeting *Notch1*. As expected, *Notch1* sgRNA cells exhibited a significant downregulation of *Notch1* target genes such as *Hes1, Fzd1* or *Tcf7l2* (Extended Data Figure 1g). Fourth, we assessed the reproducibility of our results and found that the enrichment and depletion of the top and bottom 20 sgRNAs correlated well over 8 replicates in P60 animals (Extended Data Figure 1h-i).

By employing our developed *in vivo* single-cell CRISPR strategy, we could identify 9-10 distinct cell types present in the P4 and P60 mouse skin. These included major clusters such as epidermal stem cells, suprabasal cells, hair follicle cells, sebaceous glands, melanocytes, T cells, dendritic cells, and macrophages (Figure 1d-i). The expression of classic marker genes, such as *Krt14* for epidermal stem cells, *Krt10* for suprabasal or *Cd34* for bulge stem cells showed specific expression in the corresponding clusters^11^ (Figure 1j, Extended Data Figure 2a-b). Moreover, sgRNAs were uniformly detected throughout these clusters, suggesting that our strategy enables us to infer gene function across a wide range of distinct cell types and differentiation cascades of the mouse epidermis (Figure 1i). Taken together, our established *in vivo* single-cell CRISPR strategy provides a powerful tool to study multiplexed gene function across all major epidermal cell types by coupling sgRNA representation with single-cell transcriptomics

### Cancer gene mutations result in clonal expansion

Previous ultradeep sequencing efforts conducted in physiologically normal human skin revealed somatic mutations in cancer driver genes in about a quarter of normal skin cells, with the highest burden of *Notch1/2* family, *Fat1* and *p53* mutations^3^. To determine clonal expansions in the epidermis induced by the mutations in our 150 cancer genes, we computed the total sgRNA enrichment rates compared to the initial library (T0) across the P4 and P60 time points. Consistent with findings in physiologically normal human skin, the P60 epidermis exhibited a strong enrichment of guides targeting *Fat1, Notch1, Trp53* and *Notch2* (Figure 2a, Extended Data Figure 3a, Extended Data Table 1). At the P4 stage, following the series of stratification and differentiation events during embryonic skin development, *Fat1, Notch1* and *p53* mutant clones were already expanded as the sgRNAs targeting these genes were within the top group (Extended Data Figure 3b). Between the P4 and P60 time points, while *Notch2, Notch1* and *Trp53* continued to demonstrate clonal expansion subsequent to embryonic skin development, myosin heavy chain *Myh2* and cadherin *Cdh10* emerged as the top enriched mutant genes (Extended Data Figure 3c). Additionally, we identified among the top enriched perturbations less-characterized genes in the context of tumorigenesis, such as *Ryr3* (ryanodine receptor, important for intracellular calcium release) and *Xirp2* (actin-binding protein) (Figure 2a). In contrast, sgRNAs targeting CNVs in cancer such as *p63* an essential transcription factor for epidermal development and maintenance^12^ were as expected among the most strongly depleted guides at P60 (Figure 2a). Overall, our observed clonal expansion rates faithfully recapitulate the mutational burden observed in phenotypically normal human skin^3^.

**Figure 2.**
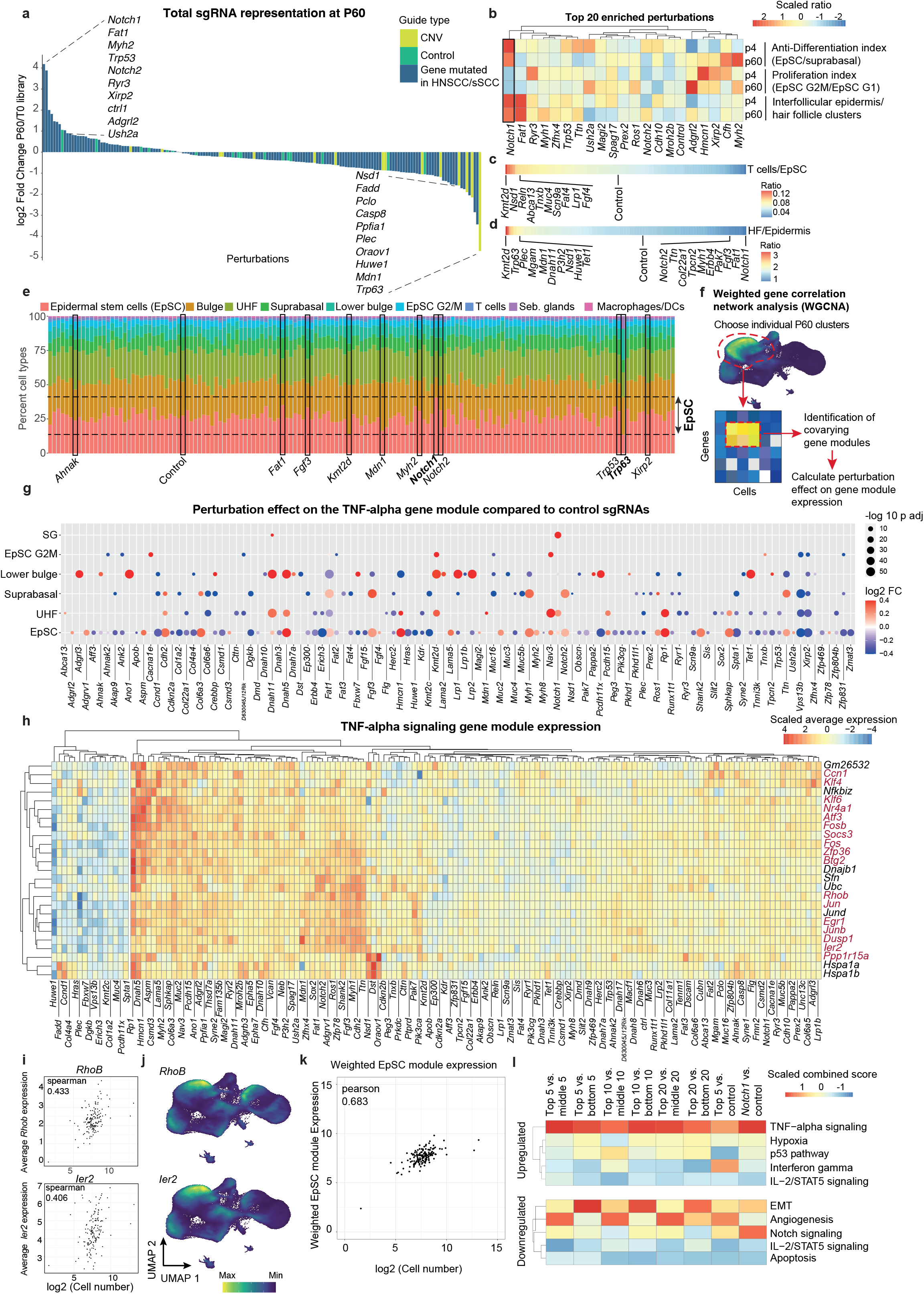
Clonal expansions in phenotypically normal epithelia uncover a shared TNF-α signaling gene module. **a**, *Notch1, Fat1* and *p53* sgRNAs are the top enriched perturbations in the P60 epidermis. The waterfall plot displays the enrichment and depletion of 150 perturbations at P60, represented as the log2 fold change between total cell numbers at P60 compared to the library T0 representation. Yellow indicates copy number variation genes, green represents control sgRNAs, and blue signifies genes mutated in head and neck (HNSCC) and skin squamous cell carcinoma (sSCC) patients. Epidermal stem cellspecific waterfall plots are shown in Extended Data Figure 3d, complete gene lists in Extended Data Table 1. Data represent the weighted average of eight P60 replicates (animals and single-cell RNA sequencing runs). **b**, Heatmap of the top 20 enriched perturbations in P60 skin. The heatmap includes the anti-differentiation index (basal/suprabasal sgRNAs ratio), proliferation index (basal G2M/basal G1 sgRNAs ratio), and epidermis index (interfollicular epidermis/hair follicle clusters 1 & 4 sgRNAs ratio). EpSC, epidermal stem cells. Control is the average of 50 non-targeting sgRNAs. **c-d**, Linear heatmaps illustrating the cell-type specificity of the perturbations. T cell-specific enrichment is calculated as the T cell/epidermal stem cell sgRNAs ratio, and hair follicle-specific enrichment is calculated as the hair follicle/epidermis sgRNAs ratio. HF, hair follicle. EpSC, epidermal stem cells. Control is the average of 50 nontargeting sgRNAs. **e**, Cell-type distribution of the 150 perturbations and the control sgRNA cells in the P60 mouse skin. The graph is alphabetically ordered and selected perturbations are indicated with black rectangles. Perturbations in *Notch1* and *Trp63*, shown in bold, result in the most enriched/depleted epidermal stem cell populations. Dashed lines indicate the minimum and maximum percentage of EpSC. **f**, Schematic outline of the weighted gene correlation network analysis (WGCNA) analysis pipeline used to identify covarying gene modules in P60 epidermal stem cells. **g**, Average perturbation effect on the TNF-α signaling gene module in the different cell types. Weighted correlation network analysis (WGCNA) has been used to describe correlation patterns among genes and for finding clusters (so-called modules) of covarying genes. WGCNA identified a major TNF-α signaling gene module present in six different cell clusters in P60 skin. Dot plots show the average log2 fold change of the genes within the WGCNA modules compared to control sgRNA cells. Dot color corresponds to log2 fold change, dot size corresponds to negative base 10 log of p adjusted. P adjusted was calculated by the Benjamini-Hochberg procedure. Module gene lists are presented in Extended Data Table 2, full WGCNA analysis in Extended Data Figure 4c. SG, sebaceous gland. EpSC, epidermal stem cells. **h**, Heatmap displaying the average expression of the 25 genes in the TNF-α epidermal stem cell module for each of the 150 perturbations. Red indicates the 17 module genes that overlap with the Gene Ontology term for TNF-α signaling. **i**, The expression of 23 out of 25 module genes positively correlated (Spearman > 0) with clonal expansions of the 150 perturbations at P60 in epidermal stem cells. Spearman correlation of two representative genes from the epidermal stem cell TNF-α module, *RhoB* and *Ier2*, show a positive correlation between clonal expansions and module gene expression. The remaining single gene correlations are displayed in Extended Data Figure 4e. **j**, Gene expression visualization by Nebulosa density plots of two module genes, *RhoB* and *Ier2*, in P60 UMAP shows dominant expression in epidermal stem cells, suprabasal cells, upper hair follicle and lower bulge. **k**, TNF-α signaling module explains a substantial portion of the variance in clonal expansion rates. A linear model was used to predict clone size with 25 module genes’ average expression in cluster 0 epidermal stem cells. Pearson correlation of the fitted linear model is 0.683. The model explains a statistically significant proportion of variance (p < 0.001). EpSC, epidermal stem cells. **l**, TNF-α signaling is the top enriched pathway in the highly enriched perturbations in the P60 skin. The top 5/10/20 perturbations were compared to either middle or bottom 5/10/20 perturbations. Transcriptomic gene set enrichment analysis was performed by Enrichr, comparing the subsets of perturbations and employing pathway analyses on the corresponding differentially upregulated and downregulated genes. Heatmap shows the scaled Enrichr combined score. Note that in *Notch1* sgRNA cells, *Notch1* signaling was also the top downregulated pathway. Pathways: MSigDB Hallmark 2020.

The interfollicular epidermis is maintained by a population of epidermal stem cells that divide stochastically, with an on-average balanced fate outcome between self-renewing daughter epidermal stem cells that remain in the basal layer and differentiating cells that migrate to the suprabasal layer to ultimately form the corneal layer^13–15^ (Figure 1f). Therefore, for clonal expansion to occur in the interfollicular epidermis, mutated epidermal stem cells need to either proliferate faster than their neighboring cells or shift the balance between self-renewal and differentiation towards the former. To further examine these two scenarios, we next analyzed the epidermal stem cell compartment. By calculating sgRNA enrichments specific to each cell type, we found that *Fat1, Notch1, Trp53* and *Notch2* were also among the top enriched mutated genes in epidermal stem cells (Extended Data Figure 3d). Next, we computed an anti-differentiation index, as defined by the guide ratios in the epidermal stem cells compared to differentiated suprabasal cells and a proliferation index, based on the number of guides in G2M cells (cluster 5) divided by the G1 epidermal stem cell guides (Figure 2b). *Notch1* sgRNA cells showed the most potent differentiation block from the top 20 expanded group, which aligns with NOTCH1’s known role in inducing suprabasal cell fate and downregulating basal fate^16,17^ (Figure 2b). However, *Notch1* knockout epidermal stem cells exhibited a low proliferation index, suggesting that *Notch1* clonal expansion is primarily driven by the differentiation block and not by increased proliferation. In contrast, epidermal stem cells with mutations in *Hmca1* or *Xirp2* showed high proliferation rates and a low differentiation block (Figure 2b). Additionally, *Notch1* and *Fat1* also exhibited the most pronounced epidermis-specific expansion rates (Figure 2b, d), highlighting cell type-specificity in clonal expansion rates and indicating that mutated *Notch1* and *Fat1* cells in the hair follicle compartment do not expand at a similar rate. This observation was further supported by assessing the total percentages of epidermal stem cells, in which *Notch1* showed the highest epidermal stem cell percentage and *p63* the lowest (Figure 2e). To further illustrate the celltype specificity of our library, we also analyzed T cell-specific sgRNA enrichment rates and found guides targeting *Kmt2d, Nsd1* and *Reln* as most strongly enriched in T cells (Figure 2c). Notably, *Kmt2d* is recurrently mutated in Sézary syndrome^18,19^, which is a leukemic and aggressive form of cutaneous T cell lymphoma with limited therapeutic options. *Reln*, on the other hand, is recurrently mutated in early T-cell precursor acute lymphoblastic leukemia^20^, indicating that our *in vivo* CRISPR strategy may also uncover genes involved in immune cell-related disorders. Together, these findings demonstrate that our singlecell CRISPR screening strategy enables cell type-specific monitoring of clonal expansion at single-cell transcriptomic resolution.

### Clonal expansion in epithelia converges on TNF-α signaling

Unlike traditional single mutation characterization, our systems-level strategy enables us to study gene programs shared by multiple cancer gene mutations. To investigate the shared gene expression changes underlying clonal expansions in epithelia, we first individually computed differential gene expression induced by the 150 perturbations in epidermal stem cells. Surprisingly, many of the top expansion-driving perturbations exhibited gene expression changes that were highly enriched for TNF-α signaling (6 out of the top 10 perturbations, Extended Data Figure 4a, Extended Data Table 2), suggesting a potential shared signaling pathway among multiple expanded cancer gene perturbations. To systematically examine shared signaling pathways, we sought to next determine how the perturbations induced by our 150 cancer gene set affects cell type-specific gene signatures. As applied in previous single-cell CRISPR screens^21–23^, weighted gene correlation network analysis (WGCNA) is a powerful tool for uncovering gene regulatory networks affected by multiple perturbations and enabling more statistical power to identify biologically relevant signaling pathways^24^. To find modules of highly correlating gene expression upon perturbation, we employed WGCNA for each of the 9 cell populations of the P60 skin (Figure 2f). In each case, we first ran a power analysis to find the power for the scale-free topology and then used all 150 perturbations to unveil gene expression changes that are common to multiple gene perturbations (Extended Data Figure 4b). We identified a total of 44 WGCNA modules across all P60 cell populations, which was strongly dominated by one major gene module that was the only module present in 6 distinct cell populations (Figure 2g, Extended Data Figure 4c, Extended Data Table 3). This major module comprised 25 genes in epidermal stem cells (Figure 2h) and gene set analysis suggested a striking enrichment for tumor necrosis factor α (TNFα) signaling, with 17 out of 25 genes overlapping with this pathway in epidermal stem cells (p adjusted = 1.21 e27, Extended Data Figure 4d, 2c). Besides this major gene cluster, we found many modules that reflected subclusters present in our major cell clusters, such as a *Dct, Tyrp1, Sox10* module in multiple clusters reflecting melanocytes within these cell populations (Extended Data Figure 4c). Another example was evident in the suprabasal compartment that encompasses spinous, granular and corneal layers, where we identified a *Lce1* (late cornified envelope) gene module, which specifically represents the late suprabasal/corneal layer within the epidermis (Extended Data Figure 4c). To summarize, our WGCNA analysis revealed various cell type-specific modules and a major module of covarying genes that displayed a pronounced enrichment for TNF-α signaling.

Clonal expansion must initiate in the epidermal stem cell compartment, as expansions in differentiated layers are ultimately shed off without long-term maintenance in epithelia. Thus, we next asked how the expression of this TNF-α gene module in epidermal stem cells correlates with expansion rates. First, testing the 25 module genes individually, we found that the expression of 23 genes positively correlated (Spearman correlation > 0) with clonal expansion of the 150 perturbations (Figure 2i-j, Extended Data Figure 4e). We then fitted a linear model^25^ to estimate how these module genes predict clonal expansions. Using the linear model, we found a Pearson correlation of 0.68 (p value = 1.4e-09), indicating that the TNF-α signaling module explains a substantial portion of the variance in clonal expansion rates (Figure 2k). Therefore, our single-cell CRISPR data suggest that early pre-cancerous mutations in cancer genes converge on a common pathway inducing a set of TNF-α signaling module genes to clonally expand in physiologically normal epithelia.

To corroborate this convergence on TNF-α signaling, we took two alternative approaches to the WGCNA analysis. First, we compared the gene expression of the top 5, 10 or 20 expanded perturbations to the corresponding average or bottom perturbed groups and performed pathway enrichment analysis. To ensure that *Notch1* and *Fat1* were not dominating the gene expression changes, we down-sampled the cell numbers to include an equal maximum number of cells per perturbation. In each of these comparisons, TNF-α signaling was the most enriched pathway among the upregulated genes, including a strong overlap with the genes of the WGCNA module (Figure 2l). Of note, although *Notch1* perturbations did not show a particularly strong impact on the 25 module genes (with 9 module genes significantly changed), *Notch1* sgRNA cells still showed TNF-α signaling as the top hallmark gene set with an additional 22 altered TNF-α pathway genes (Extended Data Figure 4f). In contrast, genes involved in epithelial to mesenchymal transition (EMT) were overall most enriched within the downregulated genes. This observation raises an interesting possibility that clonal expansions may need to be coupled to EMT downregulation to ensure that the expanding clones confine to the physiological tissue architecture of the epidermis.

In a second approach, to simplify the complex gene expression responses induced by the inactivation of our cancer genes, we also performed an unbiased clustering of transcriptomically similar cancer gene perturbations of the P60 skin (Extended Figure Data 5a). With the resulting perturbation-perturbation matrix^23^, we could identify 15 distinct clusters. For instance, *Fat1, Notch1* and *Notch2* clustered together, along with other enriched perturbations such as *Myh1* and *Fgf3*, providing additional evidence that these strongly enriched perturbations lead to similarly altered gene expression pathways. Furthermore, to visualize the perturbations specifically in epidermal stem cells, we constructed a minimum distortion embedding to place perturbations with correlated epidermal stem cell transcriptomes near each other. The embedding showed again an organization by many similarly enriched perturbations, such as clustering of the strongly enriched *Fat1, Myh1, Notch2* and *Fgf3* perturbations or a cluster with the depleted *Trp63* and *Huwe1* genes (Extended Data Figure 5b).

Collectively, despite the wide range of biological processes and functions encompassed by our cancer gene cohort, the WGCNA and tissue-wide gene expression analyses strongly suggest that clonal expansion in normal epithelia converges on a shared biological pathway mediated via a TNF-α signaling module.

### TNF-α signaling directly confers clonal expansions

Intracellular TNF-α signaling is mainly activated by binding of the TNF-α ligand to the TNF receptor 1 (TNFR1), which induces several signal transduction arms, including c-Jun and nuclear factor-κB (NF-κB) activation^26,27^. These transcription factors are responsible for diverse biological processes, including cell growth, immune and stress responses. Previous studies have demonstrated the involvement of *Tnfr1, Jnk* and *NF-κB* in epidermal proliferation^28^ and skin carcinogenesis is severely reduced in *Tnfr1* knockout mice^29^, indicating that the TNF-α signaling module could be directly mediating clonal expansions. We synthesized the effect of the different perturbations on the differential expression of the TNF-α module genes into a perturbation score by fitting a linear regression model (Figure 3a). The dynein *Dnah5*, mutated in ∼14% of HNSCC patients, the microtubule polymerization regulator *Rp1* and the hemicentin *Hmcn1*, highly enriched at P60, showed the strongest effect size (Figure 3a). In contrast, the strongly depleted guides targeting *p63* and E3 ubiquitin ligase *Huwe1* also exhibited the strongest negative effect size on the TNF-α signaling module compared to control guides (Figure 3a).

**Figure 3.**
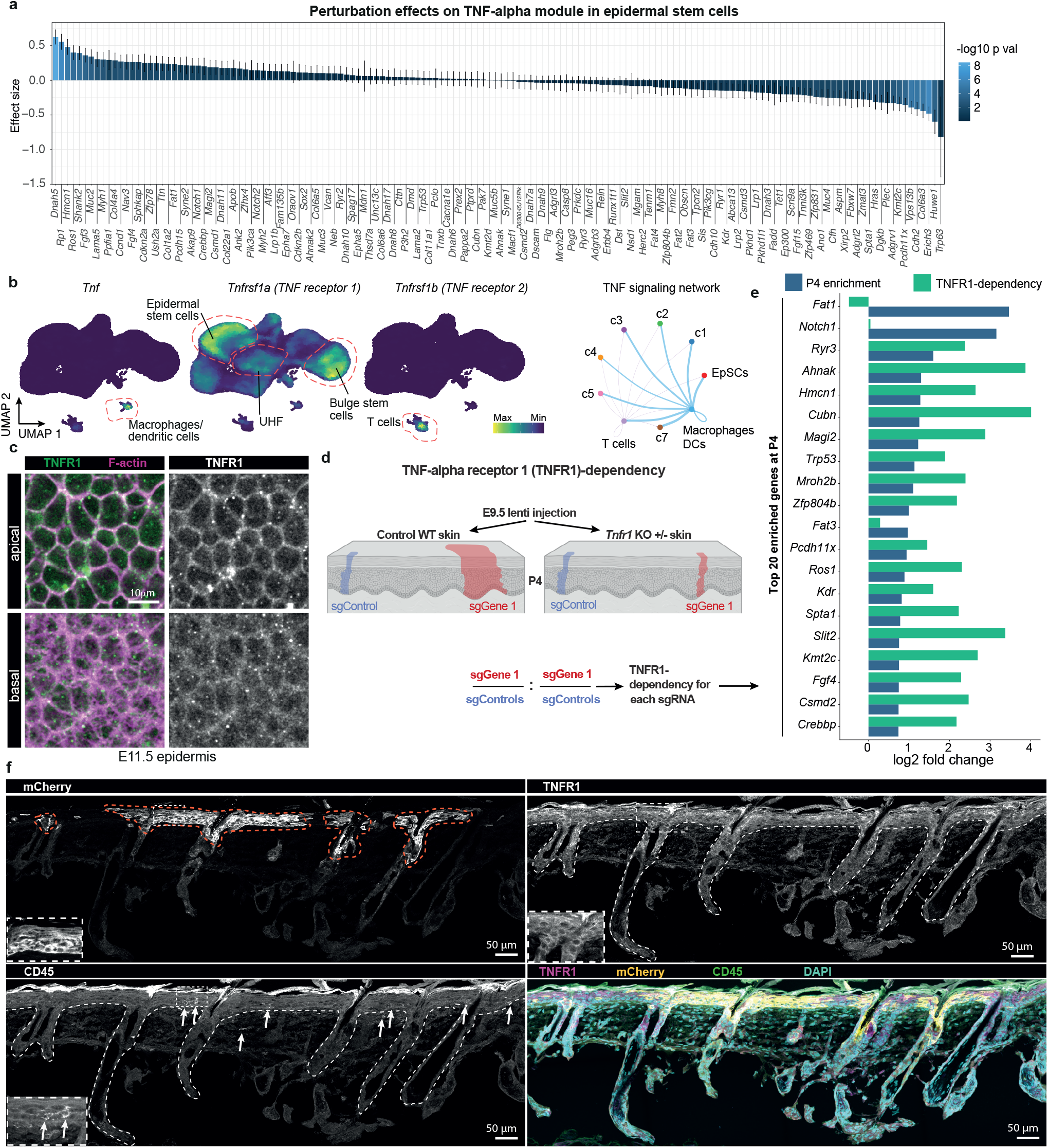
Clonal expansion in epithelia converges on TNF-α signaling. **a**, Effect size of each perturbation on the TNFmodule in epidermal stem cells compared to control sgRNAs reveals that Dnah5 and Rp1 perturbations result in the largest TNFmodule effect size. Error bars represent standard error of the linear regression. The p-values from the linear model were computed using a Wald t-distribution approximation. **b**, Gene expression visualization by Nebulosa UMAP plots highlight the cell-type-specific expression of TNF-, TNF-receptor 1 and 2 (TNFR1, TNFR2) in the P60 skin clusters. Right panel, ligand-receptor analysis by CellChat shows that the ligand TNF-is expressed mainly in macrophages/dendritic cells, which signal to the TNFR1-expressing epidermal stem cells. EpSC, epidermal stem cells. DC, dendritic cells. c-1-5 and c7 refer to the P60 clusters in Figure 1h. **c**, Whole-mount immunofluorescence shows TNFR1 cell membrane expression as early as in E11.5 embryos. Immunofluorescence images show a planar view across the basal and apical epidermis. Scale bar, 10 **μ**m. **d**, Schematic of the experimental setup to test the role of TNF-receptor 1 (TNFR1) in clonal expansions induced by the perturbations. The library with 500 sgRNAs was microinjected into E9.5 Rosa26-Cas9 or Rosa26-Cas9; Tnfr1 +/-embryos. Library representation was quantified in P4 animals by sequencing of sgRNAs. Each perturbation was normalized to the median of the 50 control sgRNAs in Rosa26-Cas9 or Rosa26-Cas9; Tnfr1 +/-animals. The ratio of normalized representation between Cas9 and Cas9; Tnfr1 +/-((WT sgRNA/WT median controls)/(Tnfr1 +/-sgRNA/Tnfr1 +/-median controls)) was used as a readout for each perturbation’s TNFR1-dependency. sgRNA/WT median controls)/(Tnfr1 +/-sgRNA/Tnfr1 +/-median controls)) was used as a readout for each perturbation’s TNFR1-dependency. **e**, TNFR1-dependency of the top 20 clonal expansions in P4 animals. All but three perturbations show a clear dependency on TNF-signaling. Data represent the average of 3 independent control and knockout animals. **f**, Immunofluorescence of mouse P60 back skin sections exhibit a mCherry-positive expanded clone, TNFR1 expression in epidermal stem cells and the presence of CD45positive immune cells in the epidermis. Insets show higher magnification of select areas. Scale bars, 50 **μ**m. xpression of the 25 genes in the TNF-α epidermal stem cell module for each of the 150 perturbations. Red indicates the 17 module genes that overlap with the Gene Ontology term for TNF-α signaling.

Next, we sought to experimentally test whether the TNF-α pathway directly mediates clonal expansion in phenotypically normal epithelia. In the P60 mouse skin, the receptor *Tnfr1* is mainly expressed in epidermal stem cells, upper hair follicles and bulge stem cells (Figure 3b). *Tnfr2* is restricted to T cells, whereas the ligand TNF is mainly secreted by macrophages (Figure 3b). Examining the ligand-receptor interactions by Cell-Chat^30^ underscored the potential main TNF-α signaling from macrophages and, to a lower level, from T cells toward epidermal stem cells (Figure 3b). To experimentally test the role of TNF-α signaling in clonal expansion, we crossed *Cas9* with *Tnfr1* +/-mice and used ultrasound-guided *in utero* microinjection to deliver our sgRNA library to target the 150 cancer genes in E9.5 embryos, resulting in equal parts of heterozygous mutant *Tnfr1* animals and control wild-type littermates (Figure 3d). We then asked which cancer gene perturbations showed *Tnfr1*-dependent clonal expansion at P4, a stage where the TNF-α signaling module is already established and we observed a robust enrichment of the perturbations such as *Notch1* and *Fat1* (Figure 3d, Extended Data Figure 3b). Of note, TNFR1 is already expressed in epidermal progenitors as early as E11.5 and located to cellular membranes (Figure 3c). Among the top 20 enriched sgRNAs at P4, all but three perturbations were highly dependent on TNFR1 for clonal expansion, indicating that TNF-α signaling plays a universal role in mediating clonal expansions in normal epithelia (Figure 3e, log2 fold changes between 1.45-4.01). Immunofluorescence analyses of epidermal P4 sections confirmed TNFR1 expression in epidermal stem cells and the presence of immune cells as a potential TNF-α source in the epidermis, including the areas of mCherry-positive clones (Figure 3f, Extended Data Figure 5c-d). Notably, exceptions to TNFR1-dependency were observed in the cases of *Notch1* and *Fat1*, two genes that showed a low proliferation index in the P4 and P60 analyses but a high anti-differentiation index (Figure 2b). This observation suggests that these two genes may rely less on TNFR1-dependent proliferation for clonal expansion and instead exploit other processes, such as a block in the differentiation cascade. Together, these results provide strong experimental evidence supporting the functional role of the TNF-α signaling module in clonal expansions in normal epithelia and establish the module as a universal response to mutations in cancer genes.

### Switch to autocrine TNF-α signaling in cancer

We next asked if and how TNF-α signaling contributes to the transition from clonal expansions to tumor initiation. *Tnfr1* knockout mice exhibit severely reduced DMBA/TPA-induced skin carcinogenesis^29^, which indicates that abrogated tumor initiation is either a consequence of inhibiting clonal expansions preceding malignant transformation or a result of a direct involvement in transformation.

To investigate the contribution of TNF-α signaling to the cellular processes underlying tumor initiation, we next induced tumors on the clonally expanded mouse skin. To this end, we injected our sgRNA library targeting the 150 cancer genes at E9.5. At the P60 stage, we induced chemical carcinogenesis with DMBA/TPA on top of the clonally expanded mouse skin for 12 weeks (Figure 4a). We collected 112 tumors (papillomas and squamous cell carcinomas) for individual tumor sgRNA capture and sorted mCherry-positive and negative tumor cells from 30 additional tumors for scRNA-seq analysis (without sgRNA capture). Of note, approximately 70% of the tumors were predominantly mCherry-positive, suggesting from a median of 2.4 % at P4 and 16.3 % infection rate at P60 -an overall positive selection of expanded clones during chemical carcinogenesis (Figure 4b).

**Figure 4.**
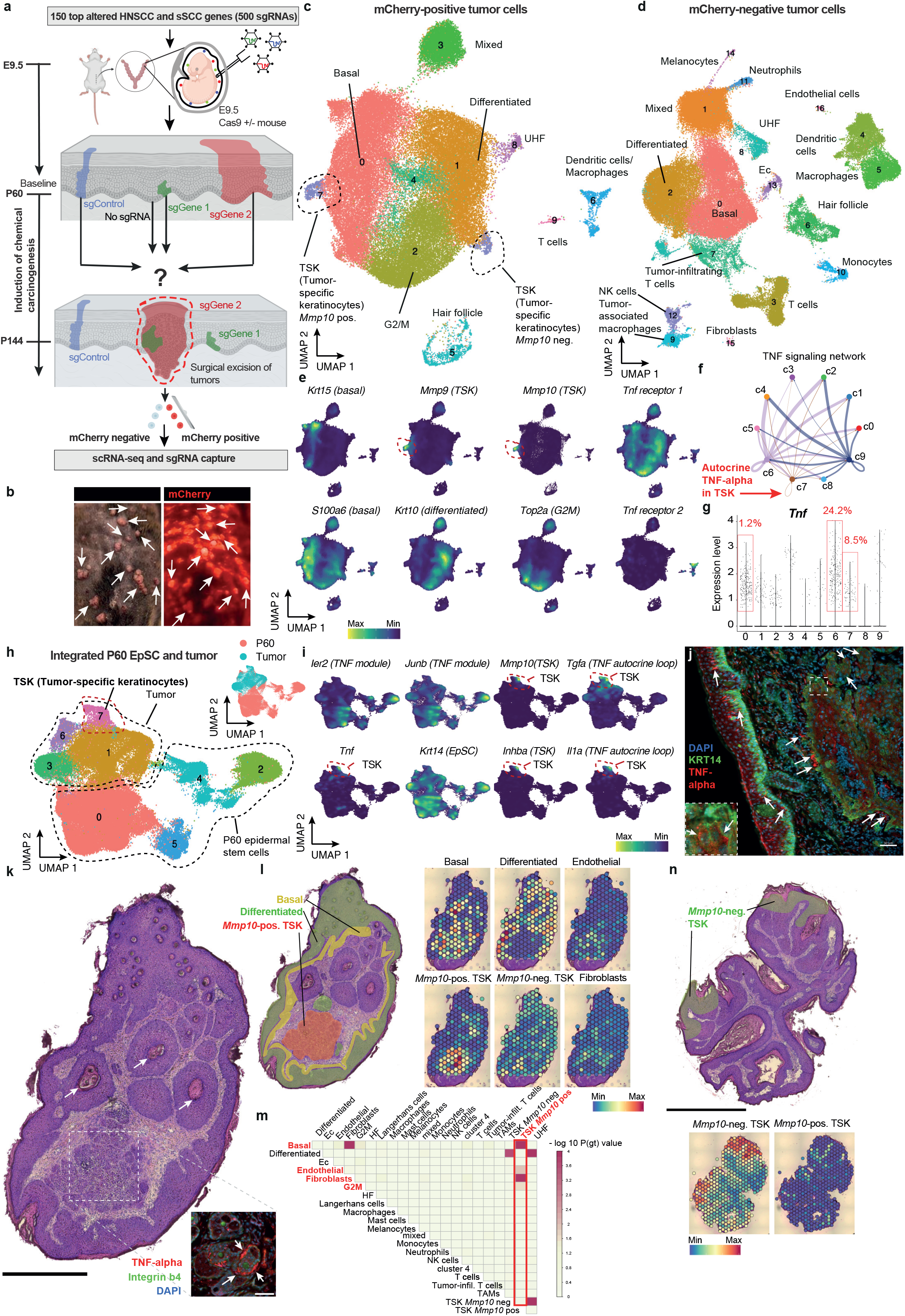
Cancer cells switch from the TNF-α module to epithelial TNF-α signaling. **a**, Schematic outline of the experimental strategy to test the role of clonal expansions in tumor initiation. The high-titer lentiviral library consisting of 500 sgRNAs is introduced into Rosa26-Cas9+/-E9.5 embryos via ultrasound-guided in utero microinjections. At P60, we induced chemical carcinogenesis by DMBA/TPA for 12 weeks and tumor single-cell suspensions were sorted for mCherry-positive and negative cells by FACS. Single-cell RNA sequencing was performed on 30 mCherry-positive tumors (from 2 animals), and another 112 tumors (from 8 animals) were collected for sgRNA amplification and sequencing of sgRNA representation. **b**, Representative macroscopic image of the mouse back skin after 12 weeks of chemical carcinogenesis by DMBA/TPA treatment show mCherry-positive and mCherry-negative tumors. Left panel, normal light. Right panel, fluorescent light. Arrows indicate individual mCherry-positive tumors. **c**, UMAP representation of single-cell RNA sequencing data from mCherry-positive tumor cells (n = 61,303 cells, from 30 tumors, 2 animals). UHF, upper hair follicle. Ec, erythrocytes. **d**, UMAP representation of single-cell RNA sequencing data from mCherry-negative tumor cells (n = 79,821 cells, from 30 tumors, 2 animals). **e**, Visualization of the tumor cell-type-specific marker gene expressions by Nebulosa density plots, highlighting the basal, differentiated, G2M and tumor-specific keratino-cyte (TSK) populations in mCherry-positive tumor UMAP. **f**, CellChat circle plot showing ligand-receptor interactions of the TNF-α signaling pathway across mCherry-positive tumor clusters, indicating potential autocrine TNF-α signaling of the tumor-specific keratinocyte (TSK) population (red arrow). **g**, Expression level and frequency of cells expressing TNF-α within each cluster of mCherry-positive tumors. h, UMAP representation of integrated P60 epidermal stem cells and mCherry-positive tumor clusters 0 and 7 using reciprocal PCA (RPCA) in Seurat. Inset shows the UMAP representation of P60 and tumor cells. **i**, Visualization of marker gene expressions by Nebulosa density plots in integrated P60 and tumor UMAP shows downregulation of TNF-α module genes Ier2 and Junb in tumors and specific expression of Mmp10 and Inhba in tumor-specific keratinocyte (TSK) populations. Tgfa and Il1a, two genes implicated in a TNF-α autocrine cascade, are also specifically expressed in the TSK population. **j**, Immunofluoresence of a squamous cell carcinoma section shows regions with TNF-α-expressing tumor cells (arrows). KRT14 serves as a basal marker (green). Inset highlights KRT14-positive TNF-α-expressing basal tumor cell. Scale bars, 50 **μ**m. **k**, Hematoxylin and Eosin (H&E) stained section of one of the tumors used for Visium spatial transcriptomics. Arrows indicate keratin pearls, a histological hallmark of squamous cell carcinomas. Box shows the Mmp10-positive tumor-specific keratinocyte (TSK) region of the tumor. The inset shows an immunofluorescence image of the TSK region with TNF-α-expressing tumor cell nests, embedded with the basal membrane marker β4-integrin. Note that the immunofluorescence inset was acquired from the same tumor and region, but from a different section. Scale bar H&E, 0.5 mm. Scale bar inset, 50 **μ**m. **l**, Visium spatial transcriptomics and spatial plots show the localization of basal, differential, Mmp10-positive tumor-specific keratinocytes (TSK), Mmp-10-negative TSK, endothelial cells and fibroblasts in the tumor, deconvoluted from the single-cell reference (see Extended Data Figure 7b-c and Methods). The left panel shows the spatial distribution of the basal, differentiated Mmp10-positive TSKs in Hematoxylin and Eosin (H&E) stained tumor section based on the histology and spatial plots. **m**, Co-occurrence analysis shows a positive correlation between the localization of Mmp10-positive TSKs, basal tumor cells, endothelia and fibroblasts, in line with Mmp10-positive TSKs residing in a fibrovascular niche. In contrast, Mmp-10-negative TSKs are positively correlating with differentiated tumor cells. P(gt), the probability of the observed co-occurrence to be greater than the expected co-occurrence. Negative correlation co-occurrence analysis is shown in Extended Data Figure 7d. HF, hair follicle. Ec, erythrocytes. Cluster 4, cluster 4 in mCherry-positive tumor. UHF, upper hair follicle. TAM, tumor-associated macrophages. **n**, H&E stained section of a second tumor used for Visium spatial transcriptomics and spatial plots predominantly show Mmp-10-negative tumor-specific keratinocytes (TSKs). Scale bar, 1 mm.

The scRNA-seq analysis of the mCherry-positive and negative tumors unveiled distinct tumor cell clusters, including basal, differentiated and cycling tumor cells (Figure 4c-e). Consistent with previous reports in human skin SCCs, we also observed a tumor-specific keratinocyte (TSK) population, located at the invasive front^31^ and expressing marker genes such as *Mmp9, Mmp10* or *Vegfa* (Figure 4c-e, Extended Data Figure 6a). Additionally, stromal cells as well as various immune cell populations were identified (Figure 4c-d, Extended Data Figure 6b). As expected, mCherry-negative tumor cells contained a higher portion of immune and stromal cells compared to mCherry-positive tumor cells (Figure 4d), reflecting the infiltration of additional untransduced immune cells during tumorigenesis.

To monitor the expression of our TNF-α module genes in the different tumor clusters, we next integrated P60 epidermal stem cells and mCherry-positive tumor cells using reciprocal PCA (RPCA) in Seurat^32^ (Figure 4h). Comparing basal tumor cells to the corresponding P60 epidermal stem cells, we observed that the large majority of TNF-α signaling module genes, including *Ier2* and *Junb*, were clearly downregulated in the different tumor clusters (Figure 4i, Extended Data Figure 6c). Gene set enrichment analysis (GSEA) comparing basal tumor cells versus P60 epidermal stem cells (control, *Notch1* or *Fat1* sgRNA) further supported the significant downregulation of TNF-α signaling and the upregulation of E2F targets and G2/M checkpoint genes, indicative of increased tumor cell proliferation (Extended Data Figure 6d). Consistent with this observation, TNF-α signaling is also strongly downregulated in human skin SCCs^33^ compared to normal skin, including an overlap with 14 downregulated TNF-α signaling module genes (Extended Data Table 4). Together, these observations imply a dual mode for the TNF-α signaling module: as the underlying signaling pathway inducing clonal expansions of cancer mutations in phenotypically normal epithelia, followed by a strong downregulation of the TNF-α module genes in early stages of tumorigenesis.

We subsequently directed our attention to the TSK cluster as these cells reside at the invasive front, hinting at their potential involvement in tumor progression^31^. TSKs were further subdivided into an *Mmp10*-positive and *Mmp10*-negative subpopulation (Figure 4c, left and right extension). Surprisingly, we observed that 8.5% of the TSK cluster cells robustly expressed endogenous TNF-α, which was a substantial increase from 1.2% of TNF-α-expressing cells in the basal tumor cluster and 0.3% in the P60 epidermal stem cells (Figure 4g). TNF-α expression overlapped with the TSK-markers such as *Mmp9/10* or *Inhba* in the integrated P60-tumor UMAP, suggesting the existence of a TNF-α-MMP9/10 axis (Figure 4i, e). Moreover, TNFα was co-expressed with transforming growth factor-α (TGF-α) and interleukin-1α (IL-1α), which were previously implicated in human epithelial cells as part of an autocrine cascade induced by TNF-α signaling^34^. This raises the interesting possibility of a switch toward an autocrine TNF-α mode during cancer progression. We confirmed TNF-α protein expression in distinct KRT14-positive basal tumor cells in squamous cell carcinomas (Figure 4j). Probing ligand-receptor interactions by CellChat further highlighted the TNF-α-TNF receptor interactions between the different tumor clusters and underscored a potential autocrine TNF-α loop within the TSK cluster (Figure 4f, Extended Data Figure 6e). To examine gene expression changes in TNF-α-expressing TSK cells, we compared TNF-α-positive vs. negative cells specifically in the TSK cluster and found that TNF-α signaling and EMT were the top upregulated pathways (Extended Data Figure 6f). Importantly, only 3 out of 26 TSK TNF-α signaling genes overlapped with the TNF-α signaling gene module, indicating that the TNF-α signaling mode in TSKs (herein referred to as autocrine TNF-α gene program) is distinct from the TNF-α module in clonal expansions. Moreover, while the TNF-α signaling module in clonal expansions was associated with downregulated EMT (Figure 2l), the autocrine TNF-α gene program acted in concert with EMT induction. Together, these findings suggest that a distinct, autocrine TNF-α signaling gene program is induced in TNF-α-producing TSKs, further promoting EMT in this subpopulation of cancer cells residing at the leading edges^31^.

To chart the spatial organization of the TSK cluster, we then performed Visium spatial transcriptomics of four additional tumors. We obtained transcriptomes from 1219 spots across 4 different sections at a median depth of 1809 UMI and 895 genes per spot (Extended Data Figure 7a, c). The spatial transcriptomics data, once deconvoluted using the single-cell tumor cell types as reference (Extended Data Figure 7b), allowed us to examine the spatial organization of *Mmp10-*positive and negative cancer cells in greater detail. Across the four tumors, *Mmp10*positive and *Mmp10*-negative TSKs were spatially segregated into distinct regions. *Mmp10-*positive TSKs resided at the invasive front of tumors, most evident in the section showing invading nests of TSKs adjacent to prominent keratin pearls (Figure 4k-l). Conversely, *Mmp10-*negative TSKs localized to differentiated regions at the surface of the tumor, distant from the invasive front (Figure 4n). Co-occurrence analysis^35^ confirmed a cellular neighborhood of *Mmp10-*positive TSKs with fibroblasts and endothelial cells, in line with a fibrovascular niche surrounding TSK cells (Figure 4m), aligning with a previous report^31^. Of note, *Mmp10-*positive TSKs correlated with G2M clusters, while *Mmp10-*negative TSKs showed a negative correlation with G2M, suggesting that *Mmp10-*positive TSKs reside in proliferative tumor areas (Figure 4m, Extended Data Figure 7d). Therefore, our spatial data indicate a subgrouping of TSKs based on their transcriptomic profile and suggest that *Mmp10-*positive TSKs represent the invasive front embedded within a fibrovascular niche.

Collectively, our findings imply a dual role for TNF-α and suggest that in the transition from clonal expansion to tumorigenesis, cancer cells strongly downregulate the TNF-α module exploited in the clonal expansion phase and instead switch to an autocrine TNF-α-MMP9/10 axis at the invasive front.

### The role of clonal expansion in tumor initiation

In human epithelia, clonal expansions do not necessarily lead to a higher tumor incidence, as evidenced by the presence of positively and negatively selected cancer gene mutations in tumors compared to normal epithelia^1,3^. Hence, we next sought to determine the systematic relationship between clonal expansions in epithelia and their role in cellular transformation. To this end, we individually sequenced the sgRNA amplicons in each of the 112 tumors and calculated the overall representation of each sgRNA in each tumor (Figure 5a, 4b). Overall, we found that *Fat1, Notch1, Ahnak, Notch2* and *p53* had the highest representation in tumors (Figure 5b). We then computed a selection score by comparing the ratio of tumor versus P60 epidermal stem cell representation for each sgRNA, normalized to the median of the control sgRNA library. This score served as a measure of each perturbation’s propensity to promote tumor initiation. Focusing on the top 40 perturbations in P60 skin and tumors, we observed that *Myh8, Prkdc, Ahnak, Kdr* and *Zfp469* showed the highest positive selection rates, indicating that these clonally expanded perturbations are prone to transformation (Figure 5c). For example, *Ahnak* is a tumor suppressor impeding cancer progression partly by controlling p53 and Wnt/bcatenin signaling^36,37^. Notably, *Kdr* and *Myh8* were found to be rather depleted at P60 (Extended Data Figure 3a). In contrast, *Zfhx4, Xirp2, Dscam, Vps13b* and *Nav3* were among the perturbations that displayed strong negative selection, suggesting that these mutated genes result in clonal expansions in phenotypically normal epithelia but suppress tumor initiation compared to control cells (Figure 5c). Our data highlight the full spectrum of possible relationships to tumor initiation, ranging from highly expanded clones that inhibit tumor initiation to depleted clones, which may display enhanced susceptibility to tumorigenesis once additional mutations are acquired.

**Figure 5.**
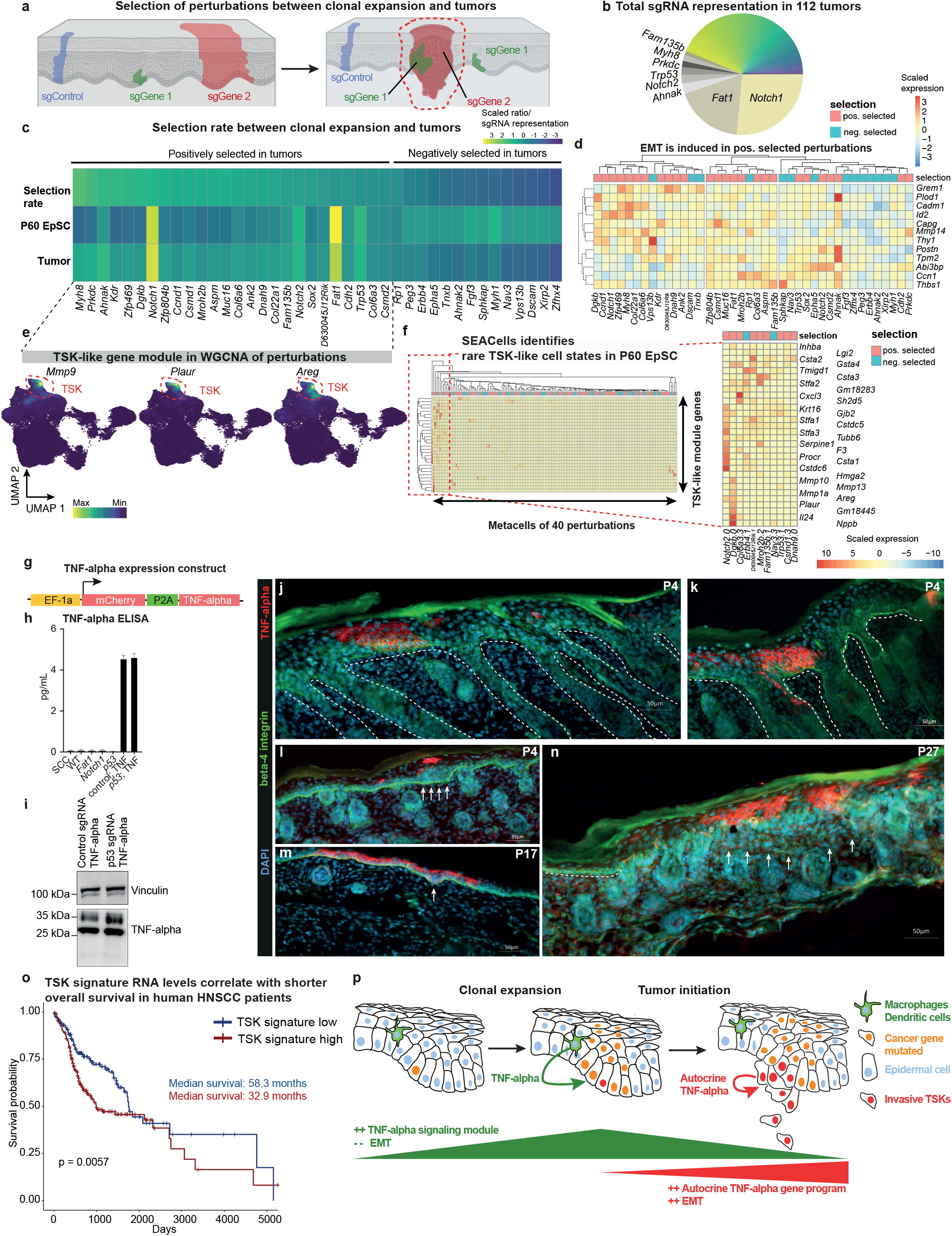
Epithelial TNF-α induces invasive properties. **a**, Schematic representation of the selection rate as a proxy for each perturbation’s predisposition for transformation. The selection rate is calculated by dividing the representation of sgRNAs in tumors following DMBA/TPA-induced chemical carcinogenesis by the representation of sgRNAs in P60 epidermal stem cells after clonal expansions. **b**, Predominance of *Notch1* and *Fat1* perturbations in tumors. Pie chart of total sgRNA representation across 112 sequenced tumors. Each tumor was individually collected and its sgRNA amplicons were individually sequenced. The total perturbation representation in tumors represents the sum of the portions in each of the 112 tumors. Each targeted gene is represented by a different color. **c**, *Myh8, Prkdc* and *Ahnak* are the top positively selected perturbations. The heatmap shows the selection score between P60 and tumors, revealing both positively and negatively selected perturbations. The selection score (sgRNA tumors/sgRNA EpSC P60) is compared to the selection score of the median of the control sgRNA library. P60 EpSC, total cell number of perturbation in P60 EpSCs. Tumor, representation of sgRNA in tumors, calculated as sum of the percentage in each of the 112 tumors. Shown are the top 20 in P60 EpSC (based on number of cells) and the top 20 perturbations in tumors, displayed as sorted according to the selection rate. The full list is in Extended Data Figure 8a. **d**, Enriched pathway and differential gene expression in positively selected perturbations. Differential gene expression between positively and negatively selected perturbations (Figure 5c) revealed epithelial-mesenchymal transition (EMT) as the top enriched pathway in positively selected perturbations. The heatmap displays the average expression of the 12 differentially expressed epithelial-mesenchymal transition (EMT) genes in epidermal stem cells, clustering positively and negatively selected perturbations from Figure 5c. **e**, Visualization of three TSK-like module gene expressions in integrated P60 tumor UMAP using Nebulosa density plots. The WGCNA analysis of the 40 perturbations (Figure 5c) in P60 epidermal stem cells uncovered a TSK-like gene module. The full list of modules is available in Extended Data Table 4, and additional Nebulosa density plots are provided in Extended Data Figure 8d. f, SEACells identifies rare cell states (metacells) that strongly induce a TSK-like gene module even in P60 clonally expanded epidermal stem cells. Heatmap shows average expression of TSK-like gene module identified by WGCNA. Boxed region is magnified on right side and shows the TSK-like module genes and their expression in the metacells. Metacells were identified by SEACells as outlined in Extended Data Figure 8c, full heatmap is shown in Extended Data Figure 8d. **g**, Schematic of the lentiviral TNF-α-expression vector used for microinjection into E9.5 embryos to ectopically express TNF-α in epithelial cells. **h**, TNF-α secretion in keratinocytes transduced with the TNF-α expression construct, as demonstrated by an Enzyme-linked Immunosorbent Assay (ELISA) of the cell supernatant. WT, wild type. SCC, squamous cell carcinoma cells. *Fat1, Notch1* and *p53* indicate sgRNA-infected knockout keratinocytes. **i**, Western blot of control sgRNA or p53 sgRNA keratinocytes transduced with lentiviral TNF-α expression construct showing the expression of the 26 kDa membrane-bound TNF-α. Vinculin, loading control. **j-k**, Immunofluorescence of P4 back skin sections depicting clones of TNF-α-expressing keratinocytes. High-titer lentivirus containing the TNF-α-expression construct was microinjected into E9.5 embryos. Green, β4-integrin, a basal membrane marker. Dashed lines indicate the basal membrane. **l-n**, Immunofluorescence of P4, P17 and P27 epidermal sections reveal regions of epidermal hyperproliferation (l), epithelial invaginations into the ear dermis (m) and breakdown of the basal membrane (n). Arrows indicate corresponding regions in l-n. Note that we did not detect any differences in TNF-α-expression plasmids containing p53 sgRNA (k, l, n) or control sgRNA (j, m) in P4-P27 animals. Scale bars, 50 **μ**m. **o**, TSK signature mRNA levels correlate with shorter overall survival in human head and neck squamous cell carcinoma (HNSCC) patients. The TSK signature was computed as average expression of the TSK genes *MMP9, MMP10, PTHLH, FEZ1, IL24, KCNMA1, INHBA, MAGEA4, NT5E, LAMC2 and SLITRK6*. The Kaplan-Meier plot shows the overall survival of HNSCC patients, stratified by upper tertile (high) and bottom tertile (low) TSK signature mRNA expression. P-value indicates a standard log-rank test. Additional Kaplan-Meier plots for TSK genes and cancer types are shown in Extended Data Figure 10. **p**, Working model illustrating the distinct TNF-α gene programs in clonal expansions in phenotypically normal tissue and tumorigenesis, resulting in a switch from the TNFα module to epithelial, autocrine TNF-α expression in invasive cancer cells.

Across all 150 perturbations, p63 emerged as the top positively selected perturbation for tumor formation (Extended Data Figure 8a), which was surprising considering the major depletion of p63 perturbations observed in the P60 skin (Figure 2a). However, beyond its established role in epidermal development, it has been documented that p63 +/mice exhibit spontaneous tumor formation, including skin squamous cell carcinomas. Notably, in mice with heterozygous mutations in both p63 and p53, a much higher incidence of squamous cell carcinomas and metastases was observed compared to mice with single p53 +/mutations. These findings imply that loss of p63 can cooperate with other mutations in tumorigenesis^38^.

To evaluate whether shared pathways confer a predisposition for transformation, we next asked if we can detect any transcriptomic differences between positively and negatively selected clones in tumor initiation. We first examined gene expression changes in the 26 positively versus the 14 negatively selected perturbations in epidermal stem cells at P60 (Figure 5c). Remarkably, our analysis revealed a significant enrichment of pathways related to EMT and angiogenesis, suggesting that positively selected perturbations already exhibit at P60 a predisposition for invasive features (Extended Data Figure 8b). Hierarchical clustering for these 12 EMT genes and 40 perturbations unveiled a robust separation between positively and negatively selected perturbations based on EMT expression (Figure 5d). Second, we employed SEACells^39^, which groups single cells into distinct cell states, so-called metacells, to enable a robust identification of rare cell states such as putative EMT-inducing epidermal stem cells. This analysis revealed select metacells strongly associated with EMT genes, in line with a rare subpopulation among the expanded epidermal stem cells capable of orchestrating EMT (Extended Data Figure 8c). Third, to find covarying genes and potential pathways shared by the positively selected perturbations, we again performed weighted gene correlation network analyses for the positive and negative tumor selection groups (Figure 5c) in P60 epidermal stem cells. We identified a specific gene module that caught our attention for several reasons.

First, the gene module contained several *bona fide* TSK markers such as *Inhba, Il24, Mmp9, Mmp10* and *Mmp13* (Extended Data Figure 8d, Figure 4e, Extended Data Table 5). Analysis of these module genes confirmed their predominant expression in the TSK cluster (Figure 5e, Extended Data Figure 8e), suggesting the presence of a “TSK-like” module. Second, our SEACells analysis revealed a cluster of cell states that showed a striking induction of the TSK-like module genes even in expanded P60 epidermal stem cells (Figure 5f). Interestingly, with 9 out of 11 metacells in this cluster, these rare cell states were predominantly present in the positively selected perturbations. Third, *Mmp3, Mmp9 and Mmp10* were significantly upregulated in TNF-α high versus low TSK cells (Extended Data Figure 6f, highlighted in red), in line with previous reports that epithelial TNF-α induces *Mmp* genes, leading to cell migration during tumor formation^40,41^. These observations suggest the presence of a subpopulation of TSK-like cells with invasive properties even in clonal expansions and indicate that epithelial TNF-α could induce the TSK-like gene program.

In summary, our systematic analyses of how clonal expansions positively or negatively contribute to tumor initiation uncovered an invasive, TSK-like gene program already present in positively selected clones and suggest that the predisposition for tumor initiation of expanded clones may be partly mediated via a TNF-α-TSK axis.

### Epithelial TNF-α expression induces invasive properties

Based on these observations, we hypothesized that the transition from clonal expansion to tumor initiation might be directly triggered by epithelial TNF-α. In such a scenario, TNF-α could promote invasive features by ECM-remodeling factors such as MMP9/10 in epidermal stem cells and raises the possibility that the predisposition to activate the epithelial TNF-α gene program in expanded clones contributes to transformation. To directly test this hypothesis *in vivo*, we designed a lentiviral TNF-α expression construct (Figure 5g).

We first confirmed ectopic TNF-α expression in keratinocytes *in vitro* and observed both secreted and membrane-bound TNF-α (Figure 5h-i). Next, we tested how TNF-α influenced cellular motility in scratch assays as a proxy for invasive properties and observed that TNF-α clearly induced cellular motility, indicating that TNF-α could confer invasive properties in keratinocytes (Extended Data Figure 9a). Moreover, epithelial TNF-α was induced by recombinant TNF-α, indicating a feedforward loop (Extended Data Figure 9b).

Having validated our construct, we then injected high-titerr lentivirus into E9.5 embryos to ectopically express TNF-α in epithelial cells and followed these mCherry-positive clones in a time course. Early time points such as P4 confirmed distinct TNF-α-expressing clones, surrounded by untransduced control epithelia, displaying clearly defined basal membranes (Figures 5j-k). However, as age progressed, TNF-α-expressing clones became associated with various abnormalities. These included epidermal hyperproliferation (Figure 5l) and epithelial invaginations into the dermis (Figure 5m). Finally, in P27 animals, we observed various examples of TNF-α-expressing clones where the basal membrane exhibited partial or complete dispersion, leading to invasive behavior of these TNF-α-expressing clones into the surrounding tissue (Figure 5n). Quantitative analysis of these phenotypes supported their increase over time (Extended Data Figure 9c). In line with our single-cell TSK analyses, *Mmp9* was induced by epithelial TNF-α expression, supporting the presence of a TNF-α-MMP axis (Extended Data Figure 9d). Together, these findings provide direct *in vivo* evidence that the epithelial TNF-α gene program is sufficient to induce invasive properties in epidermal stem cells.

As a final point, to explore the significance of the uncovered TNF-α-TSK axis in human squamous cell carcinomas, we asked whether mRNA levels of the TSK marker genes could stratify the overall survival of SCC patients. Notably, several TSK marker genes such as *Inhba* or *Nt5e* strongly correlated with shorter overall survival across different SCC types such as head and neck, lung and cervical SCC patients (Extended Data Figure 10a). TNF-α receptor 2 expression was associated with longer overall survival, while TNF-α and TNF-α receptor 1 correlated with shorter overall survival in head and neck and/or cervical SCC (Extended Data Figure 10b). Moreover, other TSK markers such as *Mmp9* demonstrated a consistent association with shorter overall survival across six different tumor types, indicating a potential widespread role of these TSK module genes (Extended Data Figure 10c). Based on the TSK marker genes, we created a TSK signature. The TSK signature mRNA levels stratified the overall survival of head and neck SCC patients, with a median survival difference of more than 2 years (Figure 5o). Taken together, these findings underscore the potential relevance of the invasive TSK signature in driving human cancer progression and position the TSK gene program as a target for cancer therapy.

## Discussion

Clonal expansions induced by cancer gene mutations can remodel entire tissues^2^. While the presence of common cancer gene mutations in phenotypically normal tissues hints at their direct involvement as cancer precursor lesions, some driver mutations have higher mutation frequencies in normal tissues compared to tumors or can even outcompete emerging new tumors^2–4^. These observations highlight the poorly understood spectrum of roles of clonal expansions in tumor initiation, ranging from strongly positively selected, tumor-promoting mutations to tumor-suppressive, negatively selected clonal expansions.

Here, we established a scalable in vivo single-cell CRISPR platform to systematically dissect tissue-wide clonal expansions of 150 cancer gene perturbations at single-cell transcriptomic resolution. Rather than focusing solely on single cancer gene mutations and their specific gene expression changes in tumorigenesis, our systems-level approach enabled us to elucidate gene programs and underlying principles shared among cohorts of cancer gene perturbations. Using this novel strategy, we provide multiple lines of evidence that a TNF-α signaling module consisting of 25 genes is a generalizable driver of clonal expansions in phenotypically normal skin. This gene module, which is primarily induced in epidermal stem cells by stromal-cell-derived TNF-α signaling, explains a substantial proportion of clonal expansion rates in epithelia for the 150 most frequently mutated cancer genes.

In contrast, during the critical transition from clonal expansion to tumor initiation, we observed a strong downregulation of the TNF-α signaling module. Instead, we identified a distinct TNF-α gene program in the TSK population. The TSK population resided at invasive tumor fronts, consistent with previous human SCC reports^31^, and switched to an autocrine TNF-α gene program leading to the expression of ECM-remodeling factors such as MMP9/10 to mediate invasion. Autocrine TNF-α in cancer cells has been reported to induce cell motility and invasions in several tumors^42–45^, raising the intriguing question of how autocrine TNF-α differs from immune cell-derived TNF-α and the molecular mechanisms underlying the activation of distinct gene programs by autocrine TNF-α. Together with our in vitro and *in vivo* findings that epithelial TNF-α is sufficient to induce invasive properties in epidermal stem cells, our data suggest that the switch to autocrine TNF-α is a key step in tumor invasion (Figure 5p).

By leveraging our in vivo experimental system, we traced the fate of expanded clones from phenotypically normal tissues to tumors and annotated their tumor selection rates as a readout for the predisposition of cancer gene mutations to undergo transformation. This enabled us to systematically examine the transcriptomic difference between positively and negatively selected perturbations. We found an enrichment of pathways related to EMT and angiogenesis in positively selected perturbations, indicating that the predisposition for tumor initiation in those clonal expansions is reflected early on by the upregulation of genes associated with invasive features. Even in phenotypically normal skin, we detected a subpopulation of TSK-like epidermal stem cells that express gene programs reminiscent of invasive tumor cells. Together, these findings support an intriguing model in which a TSK-like subpopulation in clonal expansions influences the predisposition for tumor initiation.

The discovery of two distinct TNF-α programs is highly relevant in the context of TNF-α inhibitors, which are widely used in clinics for rheumatoid arthritis and chronic inflammatory diseases and have shown some efficacy in a subset of cancer patients^26,46,47^. Our findings underscore the need for a better understanding of how TNF-α inhibitors impact the autocrine TNF-α gene program compared to stromal-derived TNF-α signaling. Exploiting specific vulnerabilities of autocrine TNF-α-expressing cancer cells, as demonstrated with SMAC-mimetics^48^, could serve as a potential therapeutic avenue. Our results suggest that the autocrine TNF-α gene program in cancer primarily induces invasive properties during tumor initiation and invasion, which could guide the development of targeted autocrine TNF-α inhibitors and help identify patient cohorts who will most likely benefit from therapy.

In summary, our study demonstrates the power of applying in vivo single-cell CRISPR to mammalian tissues, highlights the multifaceted roles of clonal expansions in epithelia and unveils a switch from a TNF-α gene module to an autocrine TNF-α gene program during tumor evolution. Given the strong correlation between TSK signature mRNA levels and shorter overall survival in human cancer, our findings provide a foundation for developing novel strategies for early cancer detection, prevention and therapy.

## Acknowledgements

We thank Felipe Quiroz, Marko Jovanovic, Ruben Barricarte, Sendoel and Moor lab members for critical input on the manuscript; Catharine Aquino, the FGCZ and Hans-Ulrich Schwarz for assistance. The project was supported by the European Research Council (ERC) under the European Union’s Horizon 2020 research and innovation programme (grant agreement No 759006), by an SNSF Professorship grant (grant number 176825), by the Swiss Cancer Research foundation (KFS-5023-02-2020-R), by the Helmut Horten Foundation and the National Center of Competence in Research (NCCR) on RNA and Disease funded by the SNSF.

## Author contributions

P.F.R. conducted the experiments and collected the data. P.F.R., A.S., S.B.S and U.G. performed data analysis and interpretation. A.K. and S.E. performed whole-mount stainings and image analysis. M.O. carried out in utero lentiviral injections and animal handling. F.V.F. assisted with lentiviral preparations and single-cell capture experiments. J.K. performed RNA-Rescue Spatial Transcriptomics on frozen tumor samples. P.F.R, A.S. and A.M. conceived the project. A.S. and A.M. supervised the project. X.F. provided input on the manuscript. P.F.R and A.S. wrote the manuscript, U.G., S.B.S and A.M. reviewed and approved the final version, with input from all the authors.

## Materials and Methods

### Selection of target genes and sgRNA design

500 sgRNAs were selected as initial targeting library. 50 out of the 500 sgRNAs were non-targeting control guide RNAs. 3 sgRNAs/gene targeted the top 150 most frequently mutated and copy-number varied (CNV) genes in head and neck and skin SCC. We choose genes that have a mutation frequency of ≥6% in head and neck SCC, ≥30% in skin SCC and ≥15% of copy number alterations in both. Mutation frequency was assessed using the cBioPortal for Cancer Genomics^50,51^ with the underlying datasets for skin^52,53^ and head and neck^54–58^ squamous cell carcinoma. The exact selection of genes and their mutation frequency can be found in Extended Data Figure 1. The overlap and differences between head and neck and skin SCC genes is graphically represented in Figure 1b (Venn diagram). Individual guides were designed with the Broad institute sgRNA design tool (sgRNA Designer: CRISPRko (broadinstitute.org)). The 3 top ranking guides per target were chosen. 50 non-targeting control sgRNAs were randomly picked from the Brie library^59^ (Addgene #73633). All 500 sgRNA sequences are listed in Supplementary Information Table 1.

### CROP-mCherry vector and library cloning

The puromycin resistance cassette in the original CROPseq-Guide-Puro vector (Addgene #86708) was replaced with a mCherry sequence, amplified from pAAVS1-NDi-CRISPRi (Gen1) (Addgene #73497) and cloned in via Pfl23II and MluI restriction sites for fluorescent-assisted cell sorting. The 500 sgRNA sequences were ordered as oligo pool from IDT, cloned in batch via Gibson assembly or via BsmbI restriction sites for single guides as previously described^5^. For all single-cell experiments, three different library batches were cloned and individually sequenced to assure accurate homogeneous sgRNA representation. Important plasmids created in this study will be made available via Addgene.

### High-titer lentivirus production

Production of vesicular stomatitis virus (VSV-G) pseudotyped lentivirus was performed by calcium phosphate transfection of Lenti-XTM 293T cells (TaKaRa Clontech, 632180) with CROP-mCherry and helper plasmids pMD2.G and psPAX2 (Addgene plasmids 12259 and 12260). 16 h posttransfection, media was changed to viral production media (D-MEM (Gibco 11965092), 1% Penicillin-Streptomycin-Glutamine (Gibco 10378016), 1% 100 mM Sodium Pyruvate(Gibco 11360070), 1% Sodium Bicarbonate 7.5% solution (Gibco 25080094), 5mM Sodium Butyrate (Sigma-Aldrich B5887)) and incubated at 37°C/7.5% CO2. Viral supernatant was collected 46 h after transfection and filtered through a 0.45 μm filter (Millipore Stericup® Quick Release Durapore®; S2HVU02RE). For in vivo lentiviral transduction, the viral supernatant was concentrated ∼2000 fold using a 100kDa MW cut-off Millipore Centricon ® 70 Plus (Merck Millipore; UFC710008). The virus was further concentrated by ultracentrifugation using the SW 55 Ti rotor for the Beckman Coulter Optima TM L-90 Ultracentrifuge at 45000 RPM / 4°C. Final viral particles were resuspended in viral resuspension buffer (20 mM Tris pH 8.0, 250 mM NaCl, 10 mM MgCl2, 5% sorbitol) and stored at -80 °C until used for titration or injection. For low titer virus, production of lentivirus was performed similarly as described above, and viral supernatant was collected and filtered using a 0.45 μm syringe filter (SARSTEDT AG & Co. KG; 83.1826) and stored at -80 °C until used.

### In vivo experiments

All animal experiments were conducted in strict accordance with the Swiss Animal Protection law and requirements of the Swiss Federal Office of Food Safety and Animal Welfare (BLV). The Animal Welfare Committee of the Canton of Zurich approved all animal protocols and experiments performed in this study (animal permits ZH074/2019, ZH196/2022).

Genetically modified mice of the strain Tg(B6J.129(Cg)Gt(ROSA)26Sortm1.1(CAG-cas9*,-EGFP)Fezh/J (denoted as “B6.Cas9”) were purchased from the Jackson Laboratory (strain #026179). B6.129S Tnfrsf1atm1ImxTnfrsf1btm1Imx (denoted as Tnfr1-ko, originally from Jackson lab #003243) were acquired through the Swiss Immunology Mouse Repository (SwImMR). Wild-type mice of the strain CD1-IGS (denoted as “CD1”) were purchased from Charles River. B6.Cas9 were crossbred either with Tnfr1-ko or CD1 to provide embryos heterozygous for the Cas9 allele, suitable for lentivirus injection at embryonic development stage E9.5.

Ultrasound-guided in utero injections were conducted as previously described^60^. In brief, females at E9.5 of gestation were anesthetized with isoflurane. Each embryo was injected with 0.5 μl of lentivirus, and up to eight embryos were injected per litter. Surgical procedures were limited to 30 min to ensure fast recovery.

All mice were housed at the Laboratory Animal Services Center (LASC) of the University of Zurich in individually-ventilated cages in a humidity- and light-controlled environment (22 °C, 45–50%, 12 h light/dark cycle), and had access to food and water ad libitum. B6.Cas9 males used for crossbreeding and pregnant CD1 females were housed individually. All other animals were group-housed. Successful infection of animals was controlled by eye through excitation with a dual fluorescent protein flashlight (NIGHTSEA, DFP-1). Positively infected animals were then euthanized by decapitation (P4) or with CO2 (P60), shaved and treated with hair removal cream if necessary.

P4 back skin was processed as previously described8, with minor modifications. After being surgically removed and scraped for fat, the back skin was washed once in cold PBS and then placed in dispase (Corning; 354235) for 35 min at 37°C, epidermis facing up, on an orbital shaker. Next, epidermis was separated from dermis with fine forceps, torn into smaller pieces and placed in 4 mL 0.25% Trypsin-EDTA (1X, Gibco; 25200056) for 15 min at 37°C with orbital shaking. Epidermis was washed with cold PBS and pipetted vigorously to achieve a single cell suspension. Suspension was then afterwards filtered through a 70 μm and a 40 μm strainer (Corning; 431750, 431751) consecutively, then centrifuged for 10 min at 1400rpm and resuspended in FACS buffer (PBS + 2 % FBS(-) + DAPI)

P60 back skin was processed as previously described^61^. In brief, back skin was harvested with surgical scissors and placed dermis-facing up on a styrofoam tray with pins. Fat and subcutis was scraped off with a scalpel. Fatfree skin was then washed, dermis-side facing down in 1xPBS. Skin was then placed in the same orientation in 0.5% Trypsin-EDTA (10X, Gibco; 15400054) and incubated at 37°C on an orbital shaker for 25 (females) or 50 min (males). Using a glass microscopy slide, skin was secured to the bottom of the dish and scraped again with a scalpel until it started to break. Excess cold PBS with 2% chelexed FBS (FBS(-)) was added to neutralize trypsin. Cell suspension was strained on ice first through a 70 μm filter, washed with an additional 15 mL of cold 1X PBS with 2% FBS(-) and then filtered through a 40 μm filter. Cell suspension was spun down at 400 x g for 10 min and resuspended in FACS buffer.

mCherry-positive cells were sorted on a BD FACSAria using a 70 μm nozzle and processed for single-cell capture with a BD Rhapsody Single-Cell Analysis System using BD Rhapsody Cartridge Kit (Cat No 633733). For P4 we loaded 7 separate cartridges with an average of 50,037 cells. For P60 we loaded 8 cartridges with an average of 64,625 cells. Reverse transcription from beads and sequencing library production was carried out according to manufacturer’s instructions (BD Biosciences, Doc ID: 210967 Rev. 1.0).

### Dial-out nested PCRs for sgRNA amplification

The sgRNA containing region has been amplified from BD Rhapsody beads in a separate nested PCR reaction. In PCR1, we used 100 μl KAPA HiFi HotStart ReadyMix (Roche; 07958935001), 6 μl forward primer (5’ ACACGACGCTCTTCCGATCT 3’, 10 μM), 6 μl reverse primer (5’TCTTGTG-GAAAGGACGA 3’ 10 μM), 12 μl Bead RT/PCR Enhancer reagent from BD Biosciences and 72 μl nuclease free water for a total volume of 200 μl. Rhapsody bead from each separate cartridge preparation were resuspended in PCR mix and aliquoted in 4x 50 μl reactions and quickly moved into a preheated PCR machine without allowing beads to settle with the following conditions: Initial denaturation at 95°C for 5 min, followed by 25 cycles of denaturation 95°C -30s, annealing 53°C - 30 s, extension 72°C - 20 s, and a final extension of 10 min at 72°C.

PCR1 product was pooled, beads were removed magnetically and amplicons were cleaned up with Agencourt AMPure XP beads (Beckman Coulter, A63881) according to manufacturer’s instructions before the second PCR. For PCR 2, 3 μl of cleaned up PCR1 was used as template along with 2 μl forward primer (5’ ACACGACGCTCTTCCGATCT 3’, 10 μM), 2 μl reverse primer (CAGACGTGTGCTCTTCCGATCTCTTGTGGAAAGGAC-GAAACA*C*C*G 3’, 10 μM), 18 μl nuclease free-water, along with 25 μl of KAPA HiFi HotStart ReadyMix for a total volume of 50 μl. PCR2 was performed under following conditions: Initial denaturation at 95°C for 3 min, followed by 10 cycles of denaturation 95°C - 30 s, annealing 60°C - 3 min, extension 72°C - 60 s, and a final extension of 5 min at 72°C. PCR2 product was cleaned up with Agencourt AMPure XP beads according to manufacturer’s instructions before the indexing PCR. Indexing PCR was performed according to BD Biosciences “mRNA Targeted Library Preparation” protocol (Doc ID: 210968 Rev. 3.0).

### Guide enrichment/depletion calculation

To calculate enrichment and depletion of sgRNAs we used guide-positive cell counts for samples at P4 and P60. At T0 we used read counts from the sequenced pre-injection library. As each Px (P60 or P4) sample received one of the three T0 library batches, we normalized each sample individually as follows. Counts at Px and T0 were first transformed to proportions over the total counts, either of each entire sample or library batch. Then, each Px sample guide proportions were normalized by dividing against its T0 proportion (that is, its initial representation in the matching pre-injection library). This yields a fold-change (of ratios). Fold-changes from triplets of guides targeting the same gene were then averaged together (and indicated with the symbol of the targeted gene), and so were the 50 control guides (grouped for a total of 17 pseudo-triplet controls: “ctrl_1” for control sgRNA from 1 to 3, “ctrl_2” for control sgRNA from 4 to 6, etc, and “ctrl_17” which comprised only control sgRNA 49 and 50). This resulted in 167 fold change values in each Px sample: 150 for the target genes and 17 for the pseudo-triplet controls. Ranks were defined for each Px sample based on these fold changes in decreasing order, therefore with the highest fold change ranking first. Ties (typically, fold changes with value zero) were assigned to the same lowest rank (see ‘min_rank’ function from R package ‘dplyr’). These ranks were used to compute the correlation of the top 20 and bottom 20 perturbations (guide triplets) among the eight P60 samples (biological replicates), as shown in Extended Data Figure 1h.

Finally, the fold changes obtained were also averaged across the same-timepoint samples (either P4 or P60). For this average, a weighted mean was used, with the weights being the total number of guide-positive cells for each sample. These results are reported in log2 scale in Figure 2a and in Extended Data Figure 1i and 3. For the P60 vs P4 guide representation (Extended Data Figure 3c), the fold changes shown in Extended Data Figure3a were divided by those in Extended Data Figure 3b (by matched perturbation identity) and re-ranked. Epidermal stem cell-specific sgRNA representations (as seen in Extended Data Figure 3d) were computed with the same approach, after subsetting the P4 or P60 datasets to include only epidermal stem cells (cluster 0, both in Figure 1e and Figure 1h).

### DMBA/TPA chemical carcinogenesis

Two-stage cutaneous chemical carcinogenesis was performed as previously described^62^. In brief, the back skin of 6-8 week old animals were shaved with electric hair clippers and treated once with 400 nmol 7,12-dimethylbenz[a]anthracene (DMBA)(Sigma-Aldrich D3254) per 100 μl (dissolved in acetone) application as an initiator mutagen. After a two-week rest period, 40 nmol 2-o-tetradecanoylphorbol-13-acetate (TPA) (Sigma-Aldrich 79346) dissolved in 100 μl 100% ethanol was applied twice a week. Tumor formation was monitored regularly by visual inspection and mice were euthanized after 12 weeks of treatment.

### Tumor preparation

Tumors and 1-2 mm of surrounding tissue from DMBA/TPA treated mice were excised and finely minced with a scalpel. Tumor pieces were submerged in 5 mL pre-warmed DMEM with 0.25% Trypsin/EDTA and 3 U/mL DNAse I at 37°C on an orbital shaker with rigorous resuspension every 5 min for a total of 30 min. Reaction was quenched with 1 ml FBS and strained through a 70 μM mesh, pelleted at 1400 rpm for 10 min with excess PBS and resuspended in 0.5 mL FACS buffer before sorting. Single-Cell capture was performed with BD Rhapsody as described above using Rhapsody Enhanced Cartridge Kit (BD Biosciences 664887). Reverse transcription from beads and sequencing library production was carried out according to manufacturer’s instructions (BD Biosciences, Doc ID: 210967 Rev. 1.0).

### Tumor preparation for sgRNA amplification

DMBA/TPA-induced tumors that were not used for either scRNA-seq or spatial analysis, we extracted genomic DNA using the Qiagen DNeasy Blood and Tissue Kit. sgRNA region amplified in a nested PCR. In PCR1, we used 12.5 μl KAPA HiFi HotStart ReadyMix (Roche, 07958935001), 0.75 μl forward primer (5’ CTTGTGGAAAGGACGAAACACCG 3’, 10μM), 0.75 μl reverse primer (5’ GTGTCTCAAGATCTAGTTACGCCAAGC 3’ 10μM), 1 μl of extracted genomic tumor DNA (with a concentration between 50-100 ng/μl) and 10 μl nuclease-free water for a total volume of 25 μl per tumor. PCR1 was performed using the following conditions: Initial denaturation at 98°C for 2:30 min, followed by 22 cycles of denaturation 98°C 20 s, annealing 62°C – 30 s, extension 72°C 30 s. PCR1 products for each tumor were individually column-purified with a GenElute™ PCR Clean-Up Kit (Sigma-Aldrich NA1020-1KT). For PCR2, 1 μl of cleaned up PCR1 was used as template along with 0.75 μl forward barcoded i5 primer (see Supplementary Information Table 1, 10 μM), 0.75 μl reverse barcoded i7 primer (see table, 10 μM), 10 μl nuclease free-water, along with 12.5 μl of KAPA HiFi HotStart ReadyMix for a total volume of 25 μl. Eight different i5 and i7 primers allow for a total of 64 different combinations. PCR2 was performed under following conditions: Initial denaturation at 95°C for 3 min, followed by 32 cycles of denaturation 98°C - 30 s, annealing 62°C - 3 min, extension 72°C – 30 s. PCR2 products for a maximum of 64 individually barcoded tumor amplifications were pooled and cleaned up with Agencourt AMPure XP beads (Beckman Coulter, A63881) according to manufacturer’s instructions before sequencing.

### Sequencing

With the exception for Whole Transcriptome Amplification Analysis (WTA)7 (sent to Novogene and sequenced on a S4 flowcell of an Illumina Novaseq), all single-cell or bulk sequencing was prepared as ready-made libraries and sequenced either on Illumina Novaseq or Nextseq 500 instruments at the Functional Genomics Center Zurich. Sequencing libraries were checked for peak size and concentration on a Bioanalyzer 2100 using DNA High Sensitivity Chip. Concentrations were additionally measured using a Qubit Fluorometer. Whole Transcriptome Amplification Analysis (WTAs) from Rhapsody Cartridges were sequenced in full SP flowcells at 1.8nM concentration with 20% PhiX and the following read configuration: Read1 60, I1 8, Read2 62 cycles. Tumors were sequenced with 10% PhiX, dual indexing i5 - 8bp, i7 - 8bp and Read1 64 cycles.

### Immunofluoresence

Back skin from P4 and P60 mice was scraped to remove fat, cut into strips, placed on whatman paper embedded in OCT (Tissue-Tek; 4583) frozen by placing in a liquid nitrogen cooled isopentane filled metal beaker, stored at - 80°C and sectioned at 10-12 μm thickness on a Leica CM1900, immobilized on Superforst glass slides, fixed for 10 min at room temperature with 4% paraformaldehyde, washed twice in 1xPBS before blocking in blocking solution (1% BSA, 1% Gelatin, 2.5% Normal goat serum, 2.5% Normal donkey serum, 0.30% TritonX-100, 1xPBS).

Primary antibodies (TNF-alpha [D2D4] XP[R] Rabbit anti-mouse monoclonal antibody, CST #11948, 1:300) and Purified Rat Anti-Mouse CD104 aka ITGB beta 4 (BD Pharmingen 553745, 1:300) were incubated over night at 4°C. After washing twice with 1xPBS, slides were incubated with secondary antibodies (Alexafluor488, Cy3, or AlexaFluor647, Jackson ImmunoResearch Laboratory; 1:500-1:1000) and 0.5μg/mL 4 ′, 6-diamidino-2-phenylindole (DAPI) at room temperature for 1 h. Sections were then washed again with 1x PBS, dried, covered with home-made mounting medium and sealed with nail polish before image acquisition. Pictures were acquired with a 20x objective on a ZEISS Axio Observer microscope controlled by the ZEN microscopy software (version 3.1).

### Whole mount immunofluorescence and antibodies

Dissected back skin from P4 and P60 animals injected with CROP-mCherry library were fixed in 4% PFA for 1h at room temperature. Following fixation, samples were permeabilized in 0.8% PBS-Triton overnight. All steps during staining were carried out at room temperature. Primary antibodies were diluted into blocking buffer (5% donkey serum, 2.5% fish gelatin, 1% BSA, 0.8% Triton in PBS) and were incubated for at least 16-20 h at room temperature. Samples were then washed for 3-4 h in 0.8% PBS-Triton, and incubated with secondary antibodies (in blocking buffer) together with DAPI (to label nuclei, 0.25 mg/mL) for at least 16-20 h. After staining, samples were extensively washed with 0.8% PBS-Triton every hour for 4-6 hours. Samples were then dehydrated in increasing concentrations of Ethanol: 30%, 50% and 70% in doubled-distilled water (with the pH adjusted to 9.0 with NaOH/HCL) for 1 h each, and finally in 100% Ethanol for 1 h. For tissue clearing, samples were transferred to Eppendorf tubes with 500 μl ethyl cinnamate (ECi, Sigma 112372) and shaken overnight at RT under dark conditions. Fresh ECi was used for mounting for imaging.

For fixed sections, back skin was cut into strips, embedded and frozen in OCT (Leica), and sectioned with a Leica cryostat (10-μm sections). Sections were permeabilized in 0.3% PBS-Triton for 15 min at room temperature, blocked in blocking buffer (5% donkey serum, 2.5% fish gelatin, 1% BSA, 0.3% Triton in PBS) for 15 min at room temperature. Primary antibodies in blocking buffer were incubated on slides overnight at 4°C, washed off with 0.3% PBS-Trition before adding secondary antibodies, with DAPI for 1 h at room temperature. After washing off the secondary, samples were mounted for imaging using ProLong™ Diamond Antifade Mountant (P36961, Invitrogen).

Antibodies used were as follows: rat anti-RFP (Chromotek, 5F8; 1:200), rabbit anti-RFP (MBL, PM005; 1:200), chicken anti-GFP (Abcam, ab13970; 1:200), rat anti-CD45-biotin (Biolegend, 103104; 1:200), goat anti-TNFR1 (R&D, AF-425-PB; 1:200). All secondary antibodies used were raised in a donkey host and were conjugated to Alexafluor488, Cy3, or AlexaFluor647 (Jackson ImmunoResearch Laboratory; 1:500-1:1000). 4′,6-diamidino-2-phenylindole (DAPI) was used to label nuclei (0.25 mg/mL).

### Whole mount microscopy

Whole-mount images were acquired using an LSM980/ Airyscan module combined with Tiling (where indicated) 40x multiimmersion objective (LD LCI Plan-Apochromat 40x/1.2 Imm Corr DIC [silicone oil, glycerol or water immersion], WD 0.41 mm) with glycerin as an immersion medium for cleared samples, or a 40x oil immersion objective (Plan-Apochromat 40x/1.4 Oil DIC, WD 0.13 mm) for sections.

### Whole mount image processing and analysis

All processing was done in Zen Blue, with a custom batch processing macro (Thomas Peterbauer). All images depicted are Maximum Intensity Projections.

### Keratinocyte culture

Newborn, primary mouse epidermal keratinocytes derived from B6-LSLCas9-eGFP were isolated as previously described^63^. In brief, isolated epidermal keratinocytes were cultured on 3T3-S2 feeder layer previously treated with Mitomycin-C in 0.05 mM Ca2+ E-media supplemented with 15% serum. After 3 passages on 3T3-S2 feeder layer, cells were cultured in 0.05 mM Ca2+ E-media, made in house as previously described14. The B6-LSL-Cas9-eGFP keratinocytes were transiently transfected with 1μg of a CRE-expressing plasmid in 6-wells using Lipofectamine 2000 (Invitrogen; 11668-027) to activate Cas9-GFP. Cells were then sorted for GFP positive signal on a BD Aria III and put back in culture using 0.05 mM Ca2+ E-media. For this project all further modified cell lines derived from the Cas9-GFP activated cell line and were grown at standard conditions, 37°C and 5% CO2. Squamous cell carcinoma cell line (SCC) was previously isolated from DMBA/TPA chemically induced skin tumors and brought into culture. Cell lines were tested for mycoplasma using the Mycoplasma PCR detection kit (Sigma; D9307).

### In vitro lentivirus infections

For lentiviral infections in culture, B6.Cas9 keratinocytes(see above) were plated in 6-well plate (Thermo Scientific Nunclon TM Delta Surface; 140675) at 1.5 × 105 cells per well and infected with 100-300μl of low-titer virus in the presence of infection mix (1/10 dilution of polybrene [10 mg ml−1 Sigma; 107689-100MG in PBS] in FBS[-]), by centrifuging plates at 1,100 g for 30 min at 37 °C in a Thermo Heraeus Megafuge 40R centrifuge. Infected cells were FACS sorted for mCherry on a BD Aria III. Viruses used for generation of knock-out keratinocytes carried the following sgRNAs for: Trp53_sgRNA.3, Fat1_sgRNA.3, Notch1_sgRNA.2, ctrl_44. Additionally, we used constructs carrying sgRNAs and TNFα cDNA: p53_sgRNA.3+TNFα, Notch1_sgRNA.2+TNFα, ctrl44-TNFα.

### ELISA

Mouse TNF-α uncoated ELISA Kit from Invitrogen (Ref 88-7324-22) was used according to manufacturer’s instructions to quantify TNF-α concentration in 2ml supernatants from confluent 6-wells of keratinocyte cultures after 24 h of incubation. Measurements of 96-well plates in technical and biological triplicates from separate wells were taken on a Tecan Infinite M1000Pro plate reader. Concentrations in supernatant were calculated according to included recombinant TNF-α standard curve.

### RT-qPCR

In vitro-infected keratinocytes and SCC cells were lysed in 6-wells with 1mL TRIzol Reagent and the RNA was purified by using the Direct-zol™ RNA Miniprep kit (Zymo Research). The procedure was performed according to the manufacturer protocol except for an additional 1 min centrifugation after the last washing step to completely remove eventual ethanol leftover. cDNA was synthesized using the Promega GoScript Reverse Transcription Mix, Oligo Protocol. In this procedure, 500-1000 ng of RNA were converted into oligo(dT)-primed first-strand cDNA. iTaq Universal SYBR Green Supermix was used according to the manufacturer’s protocol for RT-qPCR reaction in a QuantStudio 7 Flex (Applied Biosystem by Life Technologies). qRT-PCR primers can be found in Supplementary Information Table1. The delta-ct method in Quant Studio Real-Time PCR software (v1.3) was used for analysis and to calculate fold changes based on ct values.

### Western blot

Keratinocytes were cultured as described above in 6-wells, were washed with 1xPBS and lysed in protein sample buffer (100 mM Tris-HCl pH=6.8, 4% SDS, 20% glycerol, and 0.2 M DTT). The lysate was boiled for 5-10 min at 95°C and vortexed briefly to shear genomic DNA. Isolated proteins were stored at -20°C for later use or loaded directly (10-20 μg) and separated by 4-12% Bis-Tris SDS-PAGE electrophoresis (at 180 V, 300 W, 55 min) and transferred onto a nitrocellulose membrane (GE Healthcare). Membranes were blocked in 5% BSA(Sigma; A3059-100G) in TBS-Tween(0.1%) for 1 h at room temperature, incubated with primary antibodies in 3% BSA at 4°C overnight, washed with TBS-Tween. The secondary antibodies were added at room temperature in 3% BSA for 2 h. Western blots were developed with freshly mixed ECL solutions (GE Healthcare). The antibodies used are the following: Anti-TNF-α (Cell Signaling Technology, #11948), Anti-Vinculin (Abcam, ab129002). Secondary Goat Anti-Rabbit IgG HRP Linked Antibody (Cell Signaling Technology 7074S).

### Scratch Assay

Cells were seeded in an ibidi Culture-Insert 2 Well (Catalog number: 81176) according to the manufacturer’s recommendations. This insert creates a clean cell-free gap in a 6-or 12-well plate (Thermo Scientific, 150239). The assay was performed according to the manufacturer’s protocol (ibidi). Briefly, once confluency was reached, the insert was removed, and detached cells were washed away with PBS. New media (with or without 10ng/mL recombinant TNF-α (R&D Systems 410-MT) was added, and the wells were imaged every hour for 22 hours in an incubation chamber at 37°C and 5% CO2. The images were taken with a Zeiss Axio Observer controlled via the ZEN software. The data analysis was performed with an Image J plugin^64^ and default settings.

### TNF-α overexpression

Full length murine Tnf was PCR amplified from GFP-TNF-alpha plasmid (Addgene #28089) via Phusion PCR according to manufacturer’s protocol with the following primer: forward: 5’ agcctgctgaagcaggccggcgacgtggagga-gaaccccggccccatgagcacagaaagcatgatccgcgacgt 3’, reverse: 5’ ATATaac-gcgtCTAcagagcaatgactccaaag’. Primers created a 5’ P2A-site overhang with the opposing site added to the 3’ end of mCherry. mCherry-P2A-TNF was then together ligated into the CROP-seq vector via Pfl23 and MluI restriction site. Expression was tested with qPCR, western blot and immunofluoresence (see Figure 5).

### CRISPR guide efficiency

11 sgRNAs representing top, middle and bottom enrichment cohorts were selected and guide RNA efficiency for was assessed by infecting B6.Cas9 keratinocytes. Lentivirus was derived previously with low titer lentiviral preparations of the CROP-Seq-Puro vector and single guides. Cells were harvested after 10 days of puromycin selection (1μg/mL, Gibco; A11138-03). Genomic DNA was extracted using a DNeasy® Blood and Tissue Kit (QIAGEN; 69504), followed by Taq (NEB, M0273) PCR following manufacturer’s instructions. Amplicons were 800-1000 bp long and amplified by primers, designed with primer 3 listed in Supplementary Information Table 1.T7 Endonuclease assay was performed as described in the ALT-R™ genomic editing detection kit (IDT). Samples were quantified on a Bioanalyzer 2100 with a DNA high sensitivity chip. Primers for amplification of genomic target regions of guides can be found in Supplementary Information Table 1.

### Visium Spatial Gene Expression

The workflow of Mirzazadeh et al.^65^ for RNA-Rescue Spatial Transcriptomics (RRST) was followed to overcome low RNA integrity number (RIN) score. A microarray of fresh-frozen DMBA/TPA induced squamous cell carcinoma samples was cryo-sectioned with 10 μm thickness (Leica, CM3050S) and placed on the capture area of a spatial gene expression slide (10× Genomics, 1000188). Samples were stored in −80 °C before processing. The samples were fixed with 4% PFA and hematoxylin and eosin (H&E) staining was performed. The spatial libraries were then generated from the probe hybridization step (10× Genomics, 1000365) according to Visium Spatial Gene Expression Reagent Kits for FFPE (10× Genomics, User Guide CG000407 Rev D, 1000361). The resulting libraries were sequenced by the Genomics Facility Basel. The sequencing was performed in a paired-end manner with dual indexing (10x Genomics, 1000251) on a NovaSeq 6000 (Illumina) with a S4 flow cell. Libraries were then sequenced with the following cycle settings: Read1 101 cycles, i7 10 cycles, i5 10 cycles, Read2 101 cycles.

### Visium spatial expression data processing

Sequenced libraries were processed using Space Ranger software (version 2.0.1, 10x Genomics). Reads were aligned with the ‘spaceranger count’ pipeline to the pre-built mouse reference genome provided by 10x Genomics version 2020-A (comprising the STAR-indexed mm10 mouse genome and the GTF gene annotation from GENCODE version M23 (Ensembl 98)), and the probe set reference CSV file (Visium Mouse Transcriptome Probe Set v1.0) was provided for filtering of valid genes. The resulting filtered count matrix of features (i.e. genes) per spot barcodes were employed for the downstream analysis. Proportions of cell-type mixtures in each Visium spot were deconvoluted with the robust cell type decomposition (RCTD)^66^ method from the spacexr package (version 2.2.1, https://github.com/dmcable/spacexr), leveraging as single-cell reference the union of mCherry-positive (Figure 4c) and mCherry-negative (Figure 4d) tumor cells (merged dataset in Extended Data Figure 7b). Seurat R package (version 4.9.9.9041) was used to plot these cell-type deconvolution results. Spatial co-occurrence of cell-types was analyzed with the ISCHIA package^35^ (version 1.0.0.0), and resulting probabilities for positive (Pgt) or negative (Plt) co-occurrence were plotted in -log10 scale. Bioinformatic processing of the scRNA-seq data

The raw sequencing data comprising the BCL files were demultiplexed using the Illumina bcl2fastq program v2.20 with default options allowing one mismatch of the sample barcode sequences. The resulting fastq files were processed by the BD Rhapsody Pipeline version 1.9.1 hosted at the Seven Bridge’s cloud platform (https://www.sevenbridges.com/). We utilized a custom mouse reference genome based on the Gencode version 25 combined with our 500 sgRNA sequences. Since the BD pipeline uses STAR aligner^67^ in the backend, the custom STAR reference genome was generated by the genomeGenerate command in STAR. The Refined Putative Cell Calling option was disabled while running the pipeline. The output UMI counts data corrected by the BD Genomics RSEC (Recursive Substitution Error Correction) method were used for further downstream analysis.

### Processing of the PCR sgRNA dial-out data

The PCR dialouts data was processed by a custom in-house python script. Briefly, the script extracts three 9 nucleotide (nt) long cell barcodes out of possible 97 barcodes at specific locations from the Read 1, allowing one mismatch per barcode (Hamming distance of 1). It also obtains 8 nt UMI sequences from the Read 1, as provided in the BD manual. For valid barcodes, it counts the presence of sgRNAs in Read 2 by an exact match of the 20 nt sgRNA sequence. Next, we deduplicated the UMI counts per cell and obtained sgRNA UMI counts per cell. If multiple sgRNAs are detected in one cell, we only assigned the cell to a specific sgRNA if the sgRNA UMI count was higher than the quantile 0.99.

### Downstream processing of the scRNA-seq data

In order to remove the doublets, we utilized the Scrublet^68^ Python package. We removed doublets from each dataset separately with initial filtering of genes expressed in minimum 5 cells and expected doublet rate of 20% as detected by the BD Rhapsody. Next, we merged 7 and 8 scRNA-seq datasets for P4 and P60 respectively in Seurat 4^69^. The cells which had sgRNAs and were not doublets were selected for further analysis. We further filtered for cells with UMI counts > 500, UMI counts < than quantile 0.99 of total UMI counts (to remove outliers) and cells < 20% mitochondrial genes. After that, the data were processed by the standard Seurat pipeline of normalizing the data with NormalizeData function, which scales the counts to 10,000 per cell followed by natural log transform using log1p. We detected 2000 variable genes by the FindVariableGenes function. The normalized data was scaled by the ScaleData function followed by PCA analysis with the RunPCA function. We analyzed the contribution of each PC in explaining the variance of the data with the ElbowPlot function and selected top principal components for each dataset of P4, P60 and tumors. The selected principal components were used for finding nearest neighbors with the FindNeighbors function. We performed the clustering of the data with FindClusters function with modularity optimization by Louvain algorithm. Uniform Manifold Approximation and Projection (UMAP) dimensional reduction technique was used to visualize the data in two dimensions. We did not notice any batch effect in the P4 and P60 datasets in the UMAP projections. Hence no batch effect correction was performed. We obtained the marker genes for each cluster by the FindAllMarkers function and annotated the clusters with cell types based on the expression of known marker genes. The differential expression analysis of the scRNA-seq data for each perturbation was carried out using the MAST package^70^ and Wilcoxon rank-sum test, which yielded comparable results. The kernel density estimates for gene expression were inferred by the Nebulosa package.

Analysis of average expression and perturbation-perturbation matrix

To obtain an overview of the transcriptional phenotype of each perturbation, we calculated the average gene expression of cells for each perturbation in basal cells and all cells of P60. Next we selected the top 2000 most variable genes in those perturbations and computed heatmaps of correlation matrices using the pheatmap package with ward.D2 method for clustering. Moreover, we utilized the preserve_neighbors function from PyMDE package (pymde.org) for Minimum-Distortion Embedding of the data in two dimensions.

### sgRNA counts in tumor data

We sequenced over 112 tumors to detect the presence of sgRNAs in them. The raw sequencing data were demultiplexed with bcl2fastq function with corresponding i5 and i7 indexing barcodes. We counted the number of sgRNAs in the sample by the count command from the MAGECK^71^ package. The resulting sgRNA representations in tumors were used for total representation and selection scores in tumors.

### WGCNA analysis

For the WGCNA24 analysis, we downsampled the dataset to a maximum of 500 cells per perturbation using the subset function with downsample=500 option in Seurat. Next, we normalized the data with selected 2000 most variable genes with NormalizeData and FindVariableFeatures functions. The power calculation was performed by the pickSoftThreshold function from the WGCNA package for powers 1 to 30. Next, we ran WGCNA analysis with blockwiseModules function with signed network, minimum module size of 10 and mergeCutHeight value 0.15 and correlation function ‘bicor’ for robust correlations using the power estimate from the previous step. The cells with control sgRNA were not included for WGCNA analysis unless specified otherwise. WGCNA analysis was carried out for each cluster separately. To calculate the perturbation score, we ran moduleEigengenes function to obtain the module eigengenes (first principal component) of the given module. Next, we normalized the values of the first principal component using the scale function in R. Next, we performed linear modeling in R to obtain perturbation score for each perturbation over control with the formula: normalized gene score ∼ perturbation. We extracted the p-values, standard error and effect sizes from the linear model for plotting. This approach was similar to the one described in Jin et al^25^. We combined the p-values of differentially expressed module genes over control by the Fisher’s method using the metap R package and took the average of log fold change in order to compute the average perturbation effect.

### Cell-cell signaling analysis

In order to decipher cell-to-cell signaling between each cluster, we used the Cellchat^30^ package with truncated mean approach with minimum 10 cells for inferring signaling.

### Integration of scRNA-seq data

The integration of P60 cluster 0 and tumor mCherry positive clusters 0 and 7 was performed using Seurat IntegrateData function. First, each dataset was normalized and 2000 variable genes were obtained. Next SelectIntegrationFeatures was used to select features that are repeatedly variable across datasets for integration. Next, the datasets were scaled and PCA was performed with the common features. After that, we ran Seurat FindIntegrationAnchors function with RPCA algorithm utilizing 30 principal components followed by IntegrateData to integrate the datasets. We also compared the results with the Harmony^72^ algorithm, which yielded comparable results.

### SEACells analysis

To identify meta cell states in our scRNA-seq data, we employed the SEACells^73^ approach, which identifies a group of cells defined as meta cells to uncover cell states. First, we downsampled the P60 EpSC dataset to a maximum of 200 cells/sgRNA to obtain a similar number of meta cells for each perturbation. We chose ∼50 cells to define one meta cell (SEACell) and used the PCA to build the kernel. We ran the SEACells function with the option n_waypoint_eigs= no of SeaCell +1 and convergence_epsilon = 1e-5 for each perturbation separately resulting in 1-4 meta cells per perturbation. We fitted the SEACells model with a maximum of 100 iterations. Furthermore, we calculated the average normalized expression for each meta cell by the AverageExpression function in Seurat for plotting the gene expression heatmaps.

### General data analysis

The analysis was carried out using in-house Python 3.9 and R 4.1 scripts. Data wrangling was done with the pandas library in python and tidyverse library in R. Heatmaps were created using the ComplexHeatmap and pheatmap packages. Other plots were made using the ggplot2 library in R and seaborn library in Python.

### Pathway enrichment

The over-representation analysis for the pathway enrichment was performed by EnrichR31 (https://maayanlab.cloud/Enrichr), which is based on the Fisher’s exact test, and with custom R scripts utilizing the enrichR library on MSigDB Hallmark 2020 and other gene sets. We set a cut-off of FDR < 0.05 to select differentially expressed genes. We also carried out pathway enrichment analysis using the GSEA^31^ pre-ranked method using the GSEApy^74^(https://github.com/zqfang/GSEApy) which enables the analysis of up- and down-regulated genes simultaneously. This approach significantly improves the sensitivity of the gene set enrichment analysis. We performed 10,000 permutations and set an FDR cut off value of 0.05 to determine enriched gene sets.

### Survival analysis

TCGA data was obtained from UCSC Xena Toil75 (xena.ucsc.edu). Survival analysis was conducted for HNSCC and other TCGA tumor types in R utilizing the survival and survminer library. The Cox Proportional Hazards regression model was used to fit the gene expression data to survival to obtain the Hazard Ratio. Kaplan-Meier analysis was performed on samples with lower and upper tertiles (top ⅓ vs bottom ⅓ mRNA) of gene expression (tpm) and the p-values were computed by log-rank test. To obtain the association of survival with the TSK signature genes, we took the average expression of MMP9, MMP10, PTHLH, FEZ1, IL24, KCNMA1, INHBA, MAGEA4, NT5E, LAMC2 and SLITRK6. Kaplan-Meier analysis was performed with lower and upper tertiles of average expression of these genes.

### Data availability

The complete single-cell RNA sequencing and CROP-seq data for P4, P60 and tumors are available on the GEO GSE235325.

### Scripts

The main scripts used for the analyses in this paper will be available on our github repository of the Sendoel lab: https://github.com/sendoellab/single-cell_CRISPR.

---

**Extended Data Figure 1.**
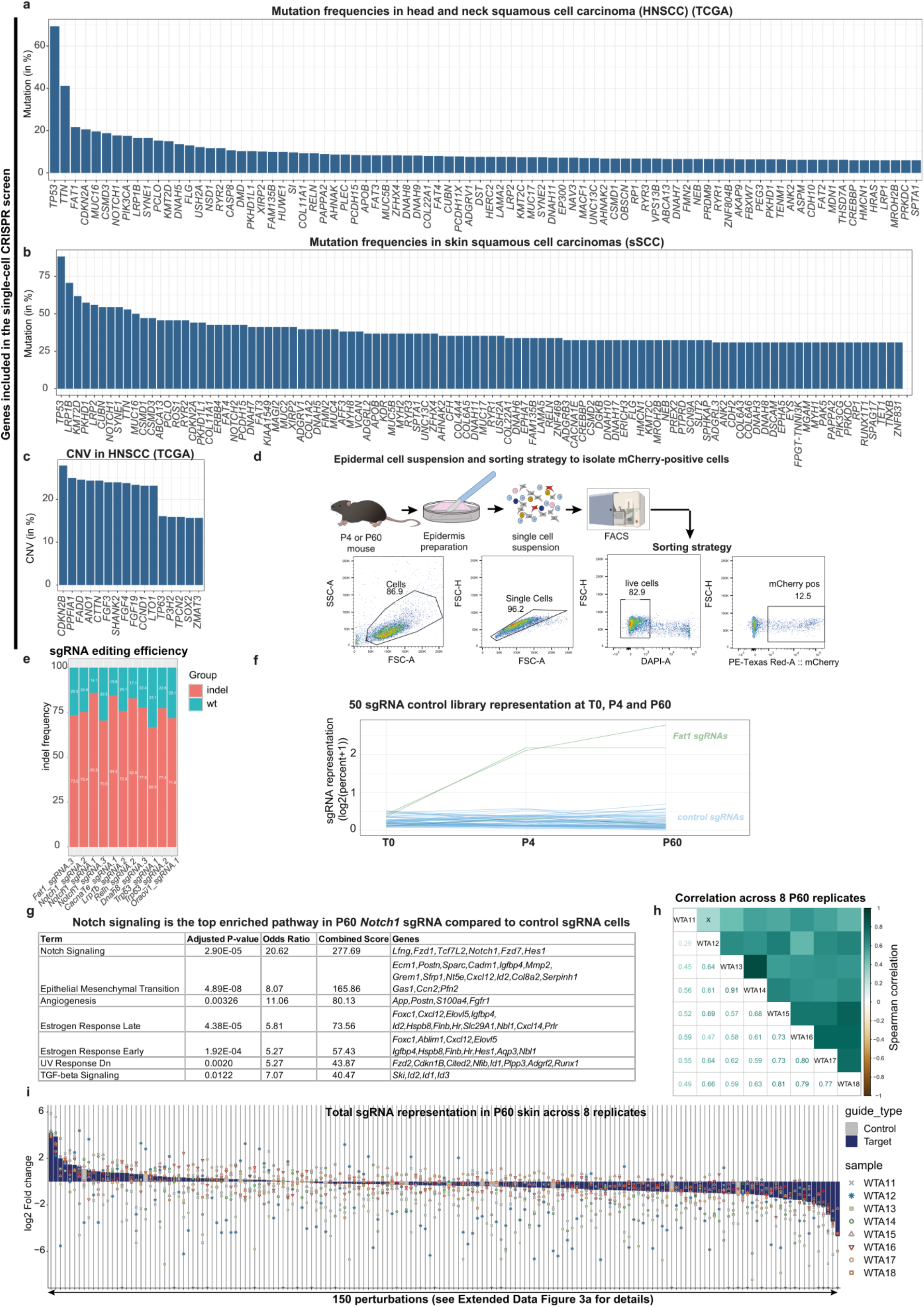
Establishing an *in vivo* single-cell CRISPR screen. a-b Perturbations included in the single-cell CRISPR library. Genes were ranked based on their mutation frequency in head and neck squamous cell carcinoma (HNSCC) and skin squamous cell carcinoma patients available from the cBioPortal^49^. c, CNVs included in the single-cell CRISPR library. Genes were ranked by their copy number variation frequency (CNV) head and neck squamous cell carcinoma (HNSCC)^49^. Besides *CKDN2B*, all CNVs are amplifications. d, Experimental workflow to prepare single-cell suspensions from back skin followed by fluorescence-assisted cell sorting (FACS) with representative gating strategy. An example preparation with 12.5 % mCherry-positive epidermal cells is depicted. Median infection rate at P4, 2.4%. Median infection rate at P60, 16.3%. e, sgRNA editing efficiency represented as the percentage of indel frequency. *Cas9*-positive keratinocytes were infected with the CROP-seq vector containing single sgRNAs. Indel frequency was measured by the Surveyor nuclease assay and corresponding DNA fragments were quantified using the Bioanalyzer. 11 sgRNAs representing top, middle and bottom enriched guides were assessed. f, Stable propagation of the control library. Non-targeting control sgRNA representation compared to the enriched sgRNA Fat1 across the T0, P4 and P60 time points. The percentage of sgRNAs relative to the total number of cells is shown. g, Gene set enrichment analysis using Enrichr of P60 *Notch1* sgRNA cells compared to control sgRNA cells reveals *Notch* signaling as the top enriched pathway, validating the bioinformatics sgRNA identification pipeline. h, P60 sgRNA enrichment and depletion correlates well across the 8 replicates, as shown by the Spearman correlation analysis of the top/bottom 20 perturbations in the P60 skin. WTA11-18 refer to the different replicates and “X” indicates FDR > 0.05. i, The enrichment and depletion patterns of total sgRNAs across the 8 replicates in the mouse P60 skin. The waterfall plot displays the enrichment and depletion of 150 perturbations at P60, represented as the log2 fold change between total cell numbers at P60 compared to the respective library T0, shown for all 8 replicates. The specific perturbation details can be found in Extended Data Figure 3a.

**Extended Data Figure 2.**
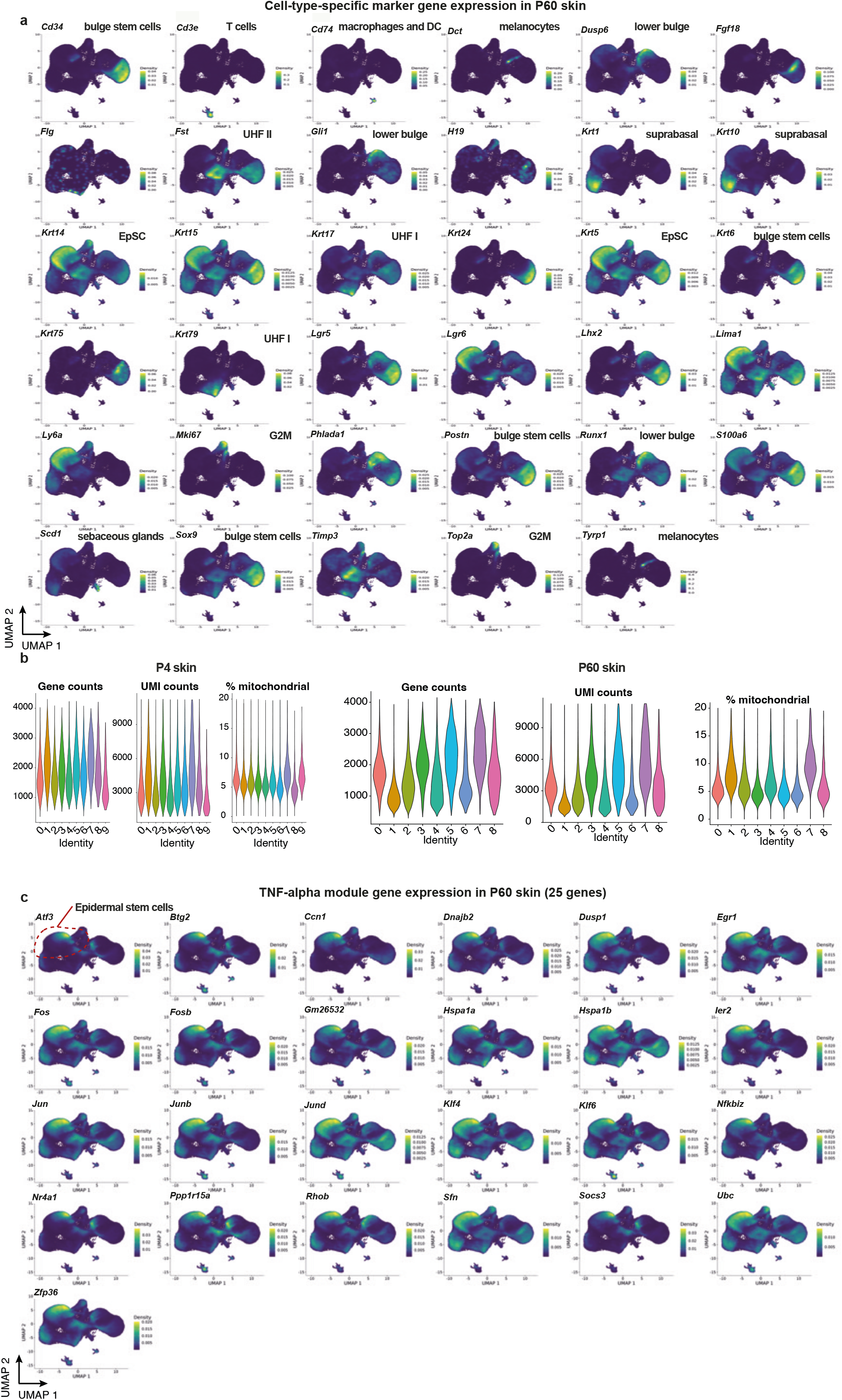
Marker gene expression in P60 UMAP. a, Visualization of cell-type specific marker gene expressions by Nebulosa density plots, highlighting their specific localization in the P60 UMAP. EpSC, epidermal stem cells. b, Characterization of the P4 and P60 single-cell RNA sequencing data sets, displaying gene counts, unique molecular identifier (UMI) counts and % mitochondrial genes as violin plots in each cluster. c, Visualization of the TNF-α signaling gene module expressions by Nebulosa density plots, highlighting their specific localization in the P60 UMAP. Red dashed circle indicates the epidermal stem cell population.

**Extended Data Figure 3.**
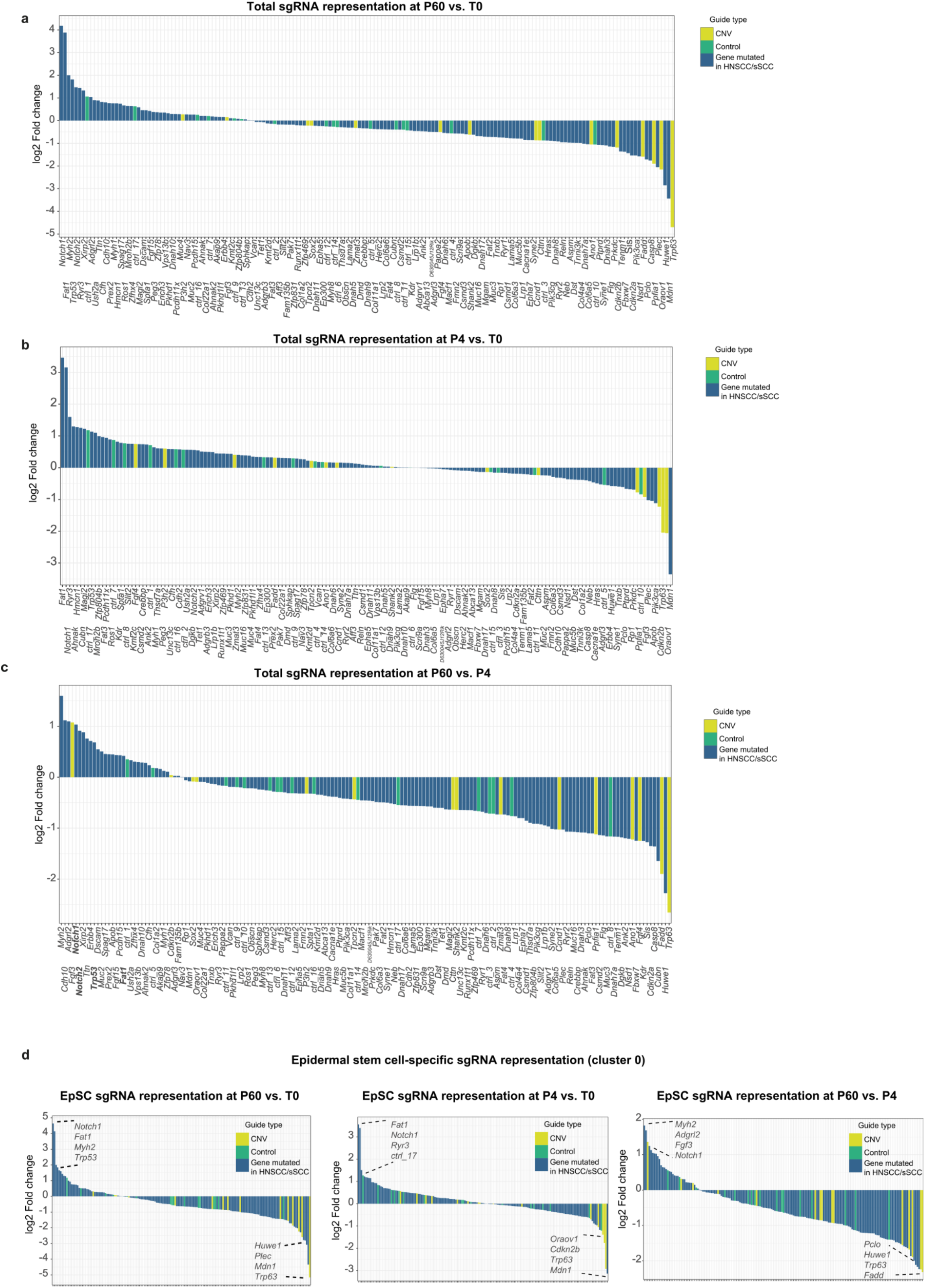
Waterfall plots showing enrichment and depletion of the 150 perturbations. a-b, The waterfall plots displays the enrichment and depletion of 150 perturbations at P60 compared to the library T0 (a), P4 compared to the library T0 (b), or P60 compared to the library P4 (c). Represented as the log2 fold change between total cell numbers. at P60 compared to the library T0 representation. Yellow indicates copy number variation genes, green represents control sgRNAs, and blue signifies genes mutated in head and neck (HNSCC) and skin squamous cell carcinoma (sSCC) patients. Data represent the weighted average of seven P4 and eight P60 replicates (7/8 singlecell RNA sequencing runs, 58 P4 animals, 10 P60 animals). d, Epidermal stem cell (EpSC)-specific enrichments. The waterfall plot displays the enrichment and depletion of 150 perturbations, represented as the log2 fold change between EpSC numbers (cluster 0) compared to T0 or P4 EpSC representation. Yellow indicates copy number variation genes, green represents control sgRNAs, and blue signifies genes mutated in head and neck (HNSCC) and skin squamous cell carcinoma (sSCC) patients. Data represent the weighted average of seven P4 and eight P60 replicates (7/8 independent single-cell RNA sequencing runs, 58 P4 animals, 10 P60 animals).

**Extended Data Figure 4.**
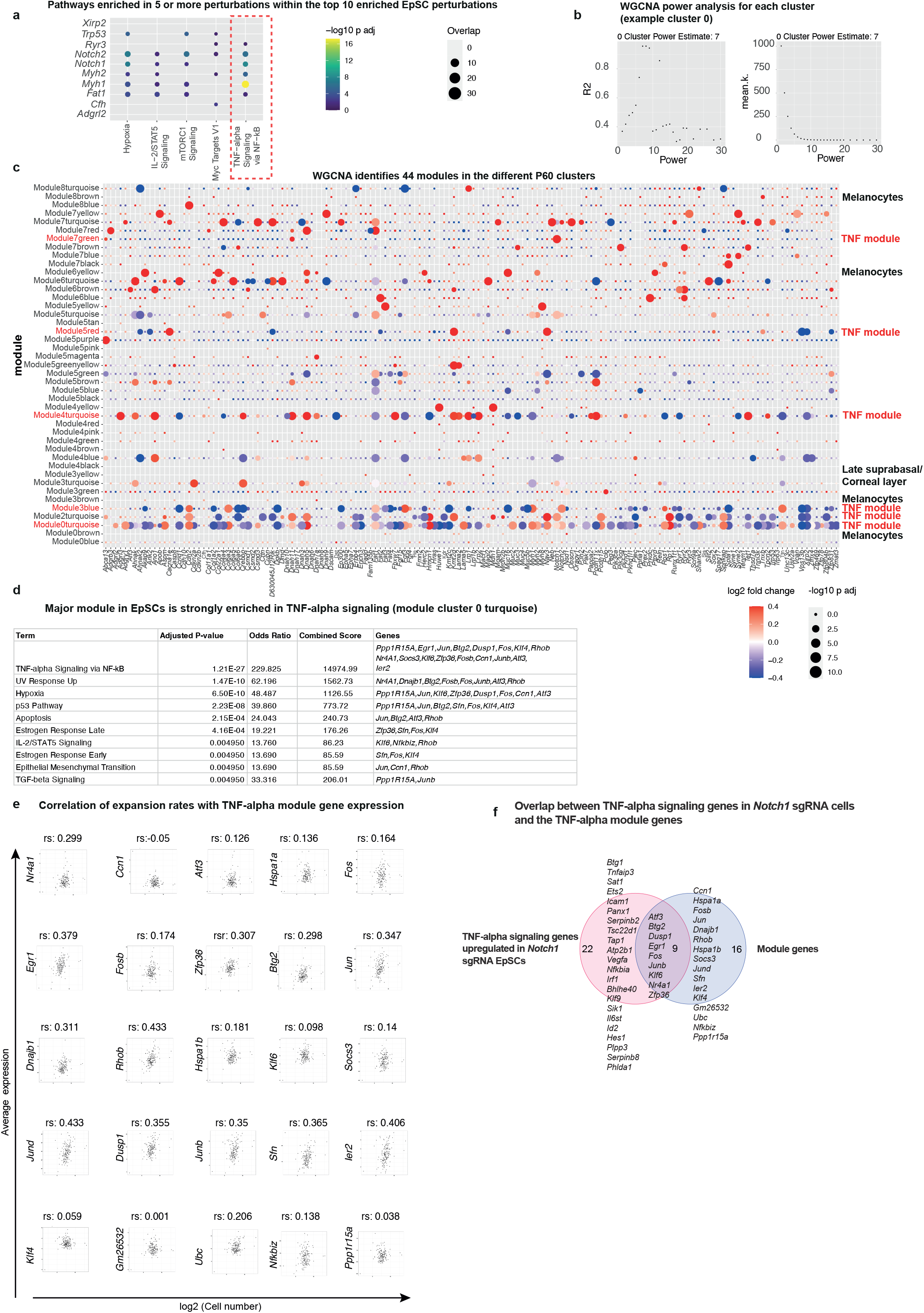
WGCNA analysis of the P60 clusters. a, TNF-α signaling is the overall most strongly enriched pathway in the top 10 expanded perturbations in P60 epidermal stem cells. Differential gene expression was computed for each of the top 10 enriched perturbations in EpSC and pathway enrichment was performed for the significantly upregulated genes. The pathway enrichment for upregulated and downregulated genes is shown in Extended Data Table 1. Dot plot shows the pathways that are in at least 5 perturbations significantly enriched. Overlap, total number of genes overlapping with pathway. b, For each cluster’s weighted gene correlation network analysis (WGCNA), a power analysis was performed to find the power for the scale-free topology and then used for all 150 perturbations to unveil gene expression changes that are common to multiple gene perturbations. An example for cluster 0, EpSC, and its power analysis is displayed. c, 44 modules were identified across all 9 P60 cell clusters. Average perturbation effect of the gene module in the different cell types. Dot plots show the average log2 fold change of the genes within the WGCNA modules, compared to control sgRNA cells. Weighted correlation network analysis (WGCNA) has been used to describe correlation patterns among genes and for finding clusters (so-called modules) of covarying genes. WGCNA identified a major TNF-α gene module, present in five different cell clusters in P60 skin and consisting of predominantly the same cohort of genes. Other gene modules represent distinct cell types within the different clusters, such as melanocytes or corneal cells within the suprabasal cells. Dot color corresponds to log2 fold change, dot size corresponds to p adjusted. Module gene lists are presented in Extended Data Table 2. P adjusted was calculated by the Benjamini-Hochberg procedure. Modules are named as Module + cluster number in P60 + color referring to the different modules in that cluster. d, The major identified gene module in epidermal stem cells (EpSC) is strongly enriched in TNF-α signaling, with 17 out of 25 genes overlapping with the pathway. Gene set enrichment analysis was performed using Enrichr. e, Spearman correlation (rs) of the expression of the 25 epidermal stem cell WGCNA module genes with clonal expansions of the 150 perturbations. The expression of 23 out of 25 module genes positively (rs > 0) correlated with expansion. f, Overlap of TNF-α signaling genes differentially expressed in P60 *Notch1* sgRNA cells compared to the TNF-α module genes. Besides 9 genes overlapping with the TNF-α module, an additional 22 genes are differentially expressed in *Notch1* sgRNA compared to control sgRNA epidermal stem cells. Pathways, MSigDB Hallmark.

**Extended Data Figure 5.**
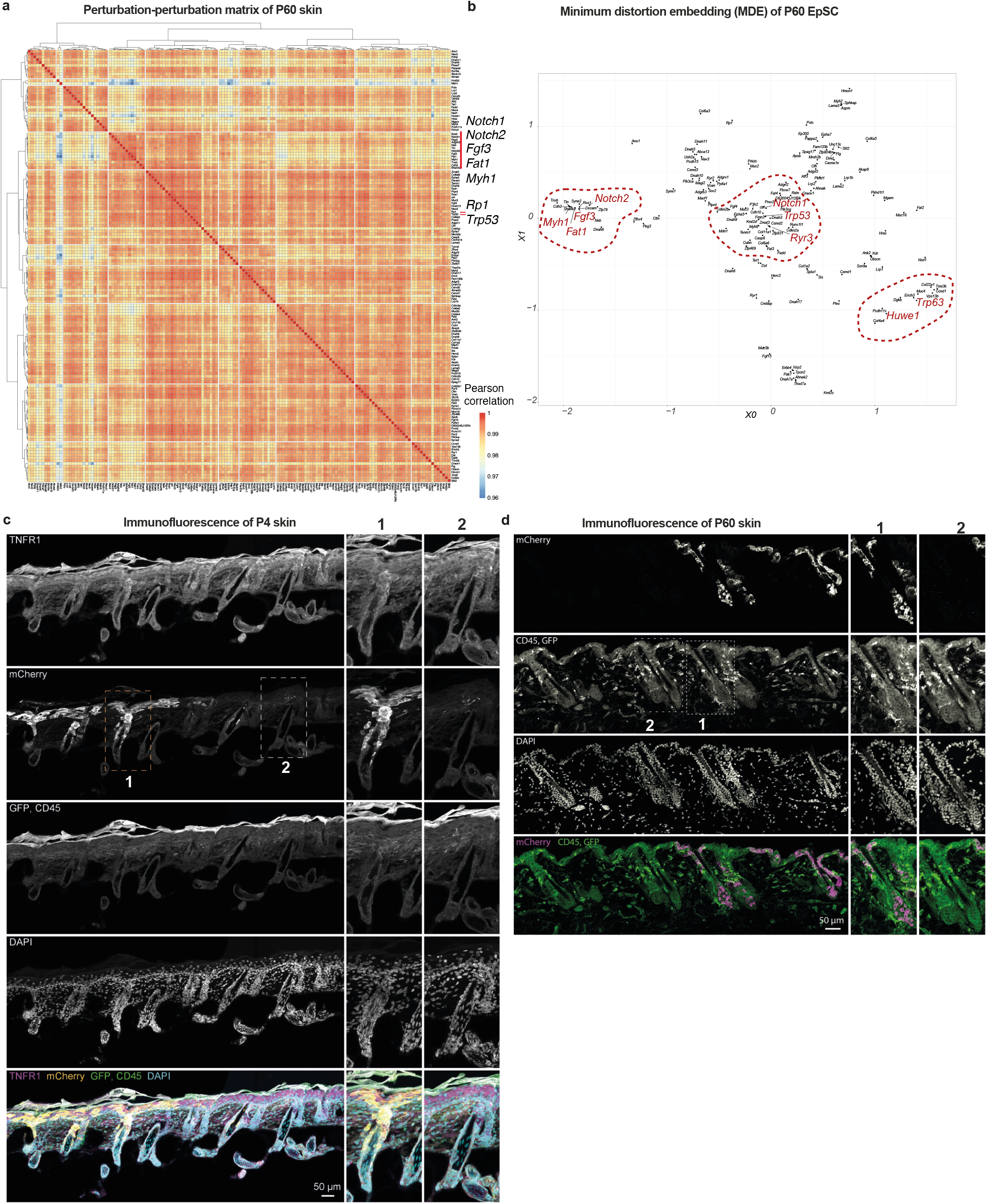
Clustering perturbations to condense expression programs. a, Clustering of perturbations to condense expression programs and phenotypes in P60 skin. P60 perturbation-perturbation matrix based on gene expression shows clustering of enriched perturbations such as *Notch1, Notch2, Fat1* and *Myh1*. b, Minimum distortion embedding (MDE) was used for the visualization and dimensionality reduction of the perturbation effect of P60 epidermal stem cells (EpSC). MDE depicting each perturbation as a dot. Some clusters are highlighted and consist of enriched and depleted perturbations in EpSCs. c-d, Immunofluorescence of mouse P4 and P60 back skin sections exhibit mCherry-positive expanded clones, TNFR1 expression in epidermal stem cells and the presence of CD45-positive immune cells in the epidermis. Insets show higher magnification of select areas. Scale bars, 50 **μ**m.

**Extended Data Figure 6.**
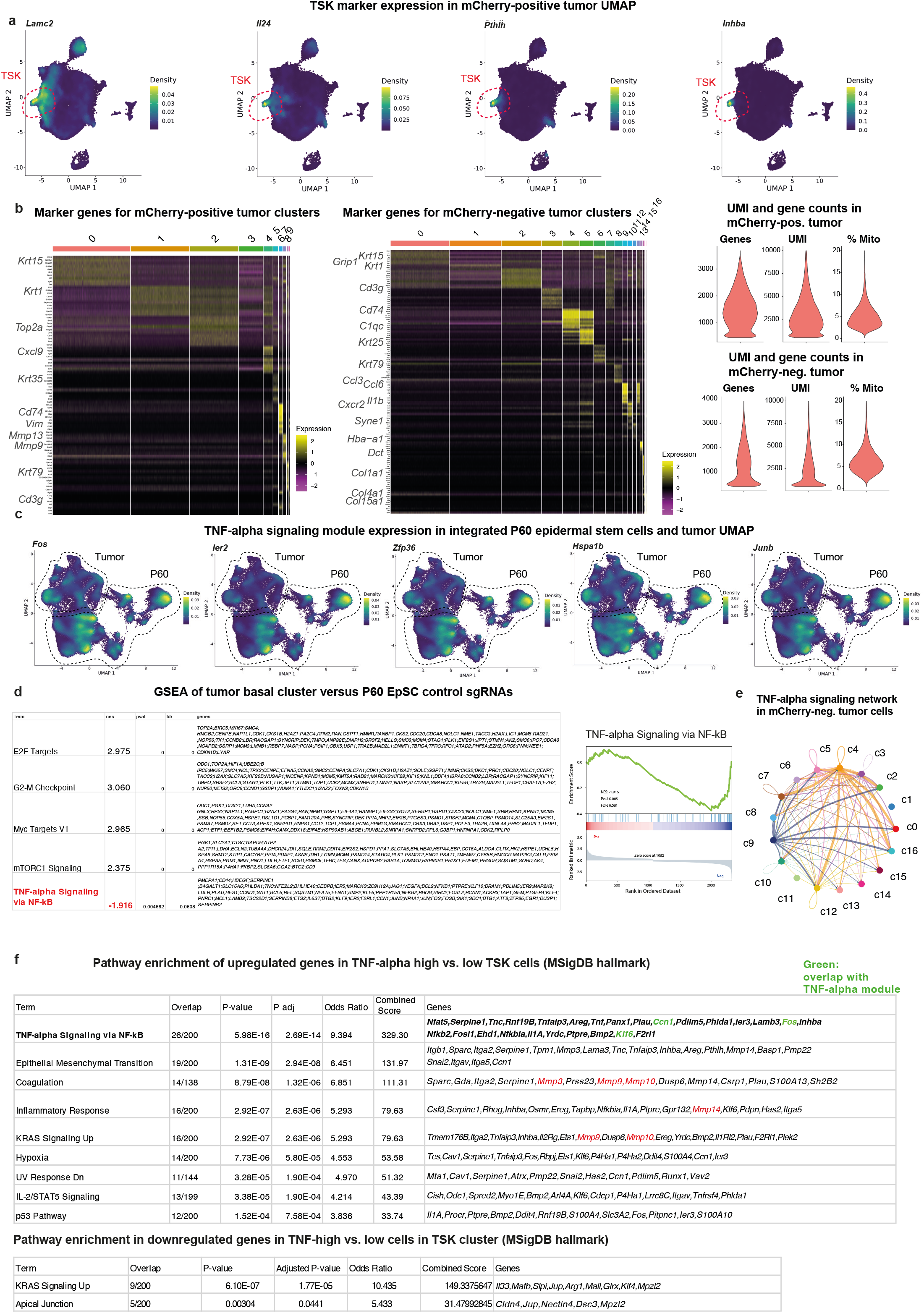
Single-cell RNA sequencing of mCherry-positive and negative tumors. a, Visualization of tumor-specific keratinocyte (TSK)-specific marker gene expressions by Nebulosa density plots, highlighting their specific localization in the mCherry-positive tumor UMAP. b, Heatmap displaying the expression profiles of the top marker genes across individual mCherry-positive and negative tumor cells (left two panels). Right panels show the quality control of the tumor single-cell RNA sequencing data sets, displaying gene counts, unique molecular identifier (UMI) counts and % mitochondrial genes as violin plots in both mCherry-positive and mCherry-negative tumor cells. c, TNF-α signaling module genes are strongly downregulated in tumors. Visualization of select TNF-α signaling gene expressions by Nebulosa density plots, highlighting their specific localization in the integrated P60 EpSC tumor UMAP. d, Gene Set Enrichment Analysis (GSEA) comparing tumor cluster 0 cells versus P60 epidermal stem cells reveals enrichment of E2F and G2M-related pathways, with TNF-α signaling identified as the top downregulated pathway. Right panel, enrichment plot for TNF-α signaling. e, CellChat circle plot showing ligand-receptor interactions of the TNF-α signaling pathway across mCherry-negative tumor clusters. f, Gene set enrichment analysis comparing TNF-α-positive tumor-specific keratinocyte (TSK) versus TNF-α-negative TSKs (cluster 7, mCherry-positive tumors) uncovers TNF-α signaling and epithelial-mesenchymal transition (EMT) as top enriched pathways in TNF-α-expressing TSKs, suggesting an autocrine TNF-α cascade associated with invasive features in TSKs. Green, only 3 genes overlap with the TNF-α module, suggesting the presence of a distinct TNF-α gene program. Red, *Mmp* genes.

**Extended Data Figure 7.**
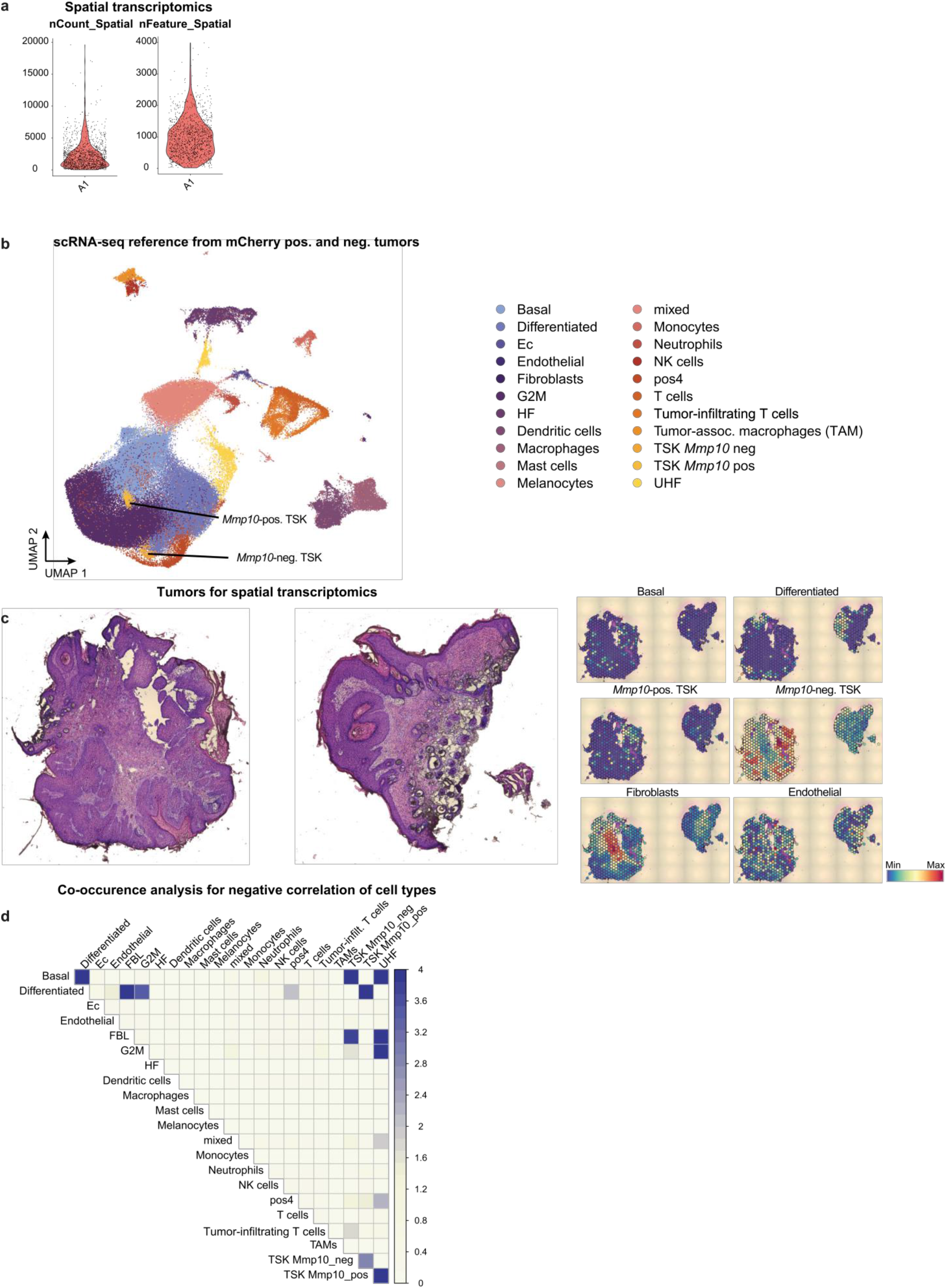
Spatial transcriptomics of tumors. a, Characterization of the tumor spatial transcriptomics datasets. Violin plots displaying gene counts (nFeature) and unique molecular identifier (UMI) counts (nCount) of the 4 processed tumor sections. b, UMAP of the single-cell reference employed for the cell-type deconvolution of the Visium spots (see Methods section). Obtained by merging the scRNAseq datasets of mCherry-positive and mCherry-negative tumors (datasets from Figure 4c-d). Ec, erythrocytes. HF, hair follicle. Pos4, cluster 4 in mCherry-positive UMAP. UHF, upper hair follicle. c, Hematoxylin and Eosin (H&E) stained images of two additional sections utilized for Visium spatial transcriptomics. A total of four sections were processed to obtain spatial transcriptomics data. Spatial plots highlight the basal and differentiated tumor areas, as well as *Mmp10*-positive and negative tumor-specific keratinocytes (TSK), fibroblasts and endothelial cells. d, Co-occurrence analysis demonstrating negative correlations between different cell types within the tumors. The probability (P(lt)) indicates the likelihood of the observed co-occurrence being less than the expected co-occurrence. Ec, erythrocytes. HF, hair follicle. Pos4, cluster 4 in mCherry-positive UMAP. FBL, fibroblasts. TAM, tumor-associated macrophages. UHF, upper hair follicle.

**Extended Data Figure 8.**
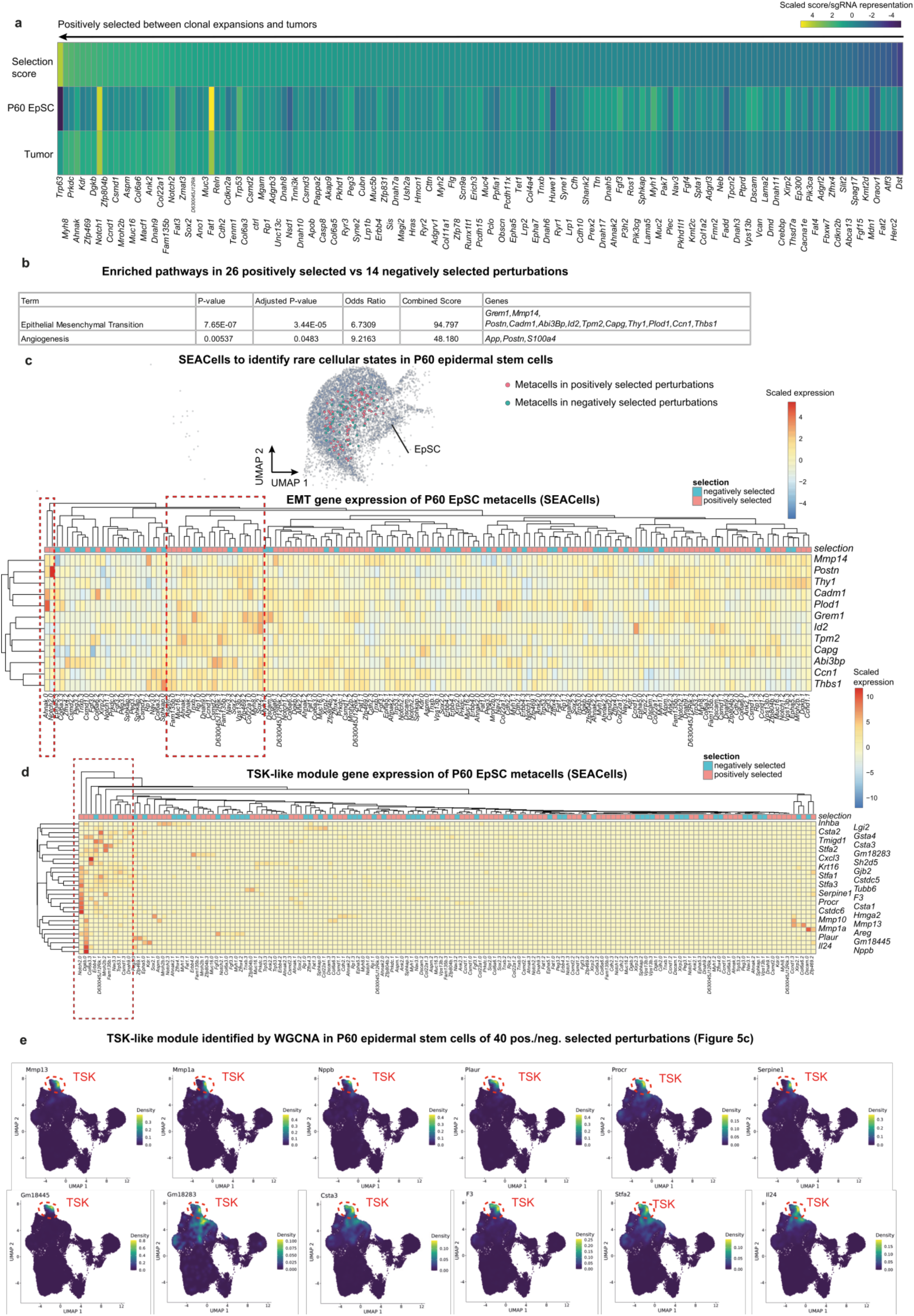
SEACells identifies rare cell states with invasive properties. a, The heatmap shows the selection score for all 150 perturbations between P60 and tumors, revealing both positively and negatively selected perturbations during tumor initiation. Because of the n = 112 tumors, we focused our main analysis on the top 20 in P60 EpSC and the top 20 in tumors to ensure the robustness of selection score, as presented in Figure 5c. The selection score (sgRNA tumor/sgRNA EpSC P60) is compared to the selection score of the median of the control sgRNA library. P60 EpSC, total cell number of perturbation in P60 EpSCs. Tumor, representation of sgRNA in tumors, calculated as sum of the percentage in each of the 112 tumors. The selection score is compared to the selection rate of the median of the control sgRNA library. EpSC, epidermal stem cells. b, Gene set enrichment analysis (GSEA) of differentially expressed genes between 26 positively selected perturbations and 14 negatively selected perturbations (as shown in Figure 5c) uncovers significant enrichments in genes associated with epithelial-mesenchymal transition (EMT) and angiogenesis. c, Identification of rare cell states in P60 epidermal stem cells by single-cell aggregation of cell states (SEACells)^39^. SEACells algorithm was applied to identify metacells in positively and negatively selected perturbations, upper panel. Positively selected and negatively selected metacells are highlighted in blue and red large cells. Lower panel, heatmap shows average gene expression of epithelial-mesenchymal transition (EMT) genes that were differentially expressed between positively and negatively selected perturbations metacells (Extended Data Figure 8b). The number next to the perturbation refers to the metacell number. Boxes highlight clusters of metacells (rare cell types) expressing high levels of EMT genes, in line with a small subpopulation with invasive features. d, Rare cell states induce TSK-like gene module even in P60 clonally expanded epidermal stem cells. Heatmap shows scaled average expression of the TSK-like module genes identified by WGCNA. Metacells were identified by SEACells as outlined in Extended Data Figure 8c. The number next to the perturbation refers to the metacell number. A closeup of the red dashed region is provided in Figure 5d. e, Visualization of the TSK-like module gene expressions by Nebulosa density plots, highlighting their specific localization in the TSK cluster in the integrated P60 tumor UMAP.

**Extended Data Figure 9.**
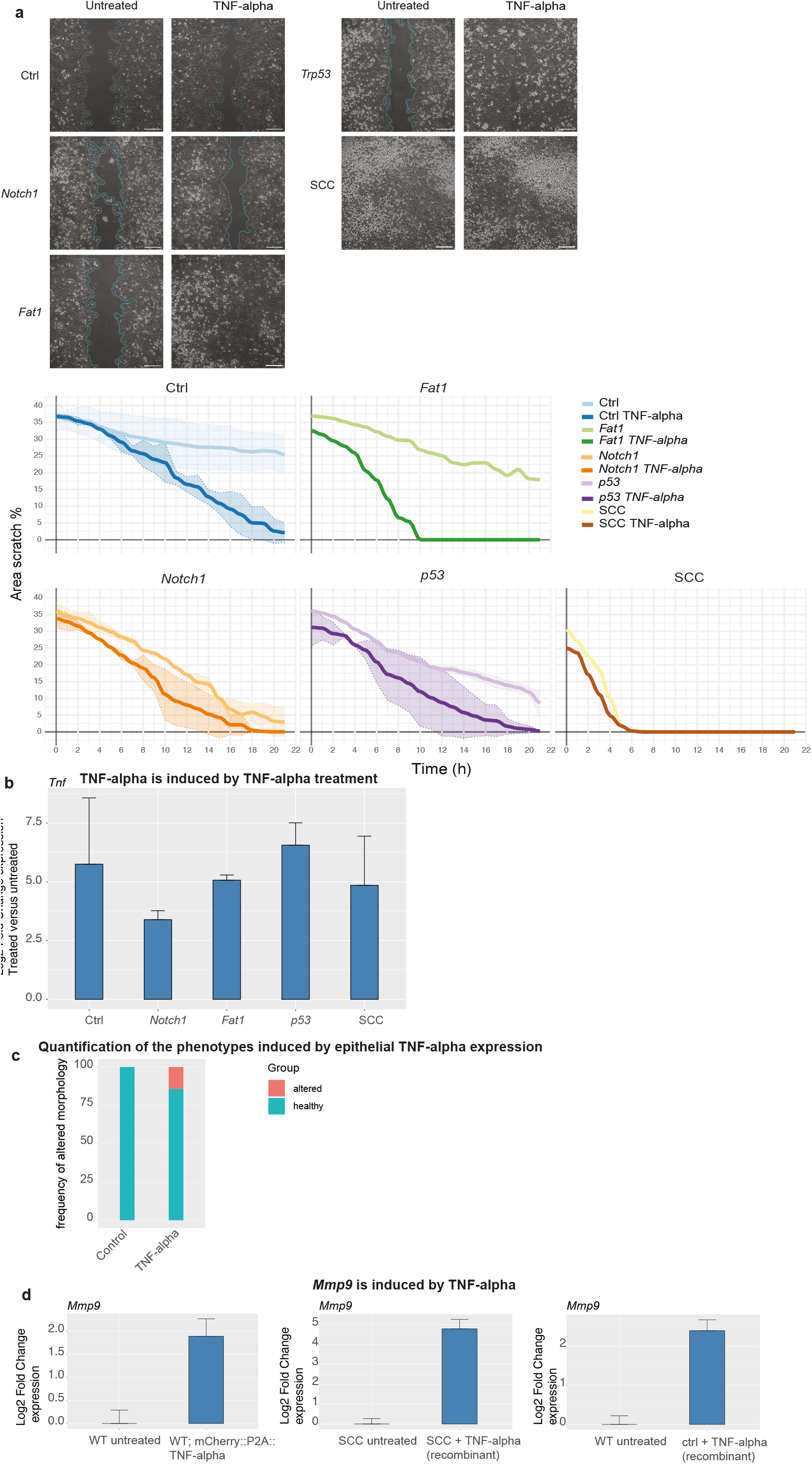
TNF-α induces motility in keratinocytes. a, *In vitro* scratch-wound assay shows that TNF-α can induce cellular motility in keratinocytes. Different perturbations in keratinocytes isolated from *Rosa26-Cas9* mice (and infected and selected with *Notch1, Fat1* and *p53* sgRNAs) and squamous cell carcinoma cells (SCC) were treated with recombinant murine TNF-α and wound closure was tracked over time. Scale bar, 200 **μ**m. b, TNF-α treatment induces epithelial TNF-α expression. Quantitative reverse transcription PCR was performed for treated and untreated keratinocytes and TNF-α induction was calculated. c, Quantification of the TNF-α expression phenotype was performed using immunofluorescence staining of P4, P17, and P27 epidermal sections obtained from animals injected with a TNF-α overexpression construct. Infected regions were identified based on TNF-α staining, while adjacent negative regions served as controls. Quantified as altered were regions that displayed hyperproliferation of epidermal stem cells, epithelial invaginations or breakdown of the basal membrane. N = 42 TNF-α-positive regions and n = 42 control regions. d, Epithelial TNF-α expression or recombinant murine TNF-α treatment induces epithelial *Mmp9* mRNA. Quantitative reverse transcription PCR for *Mmp9* was performed for treated/infected and untreated keratinocytes. The lentiviral construct TNF-α expression is displayed in Figure 5f.

**Extended Data Figure 10.**
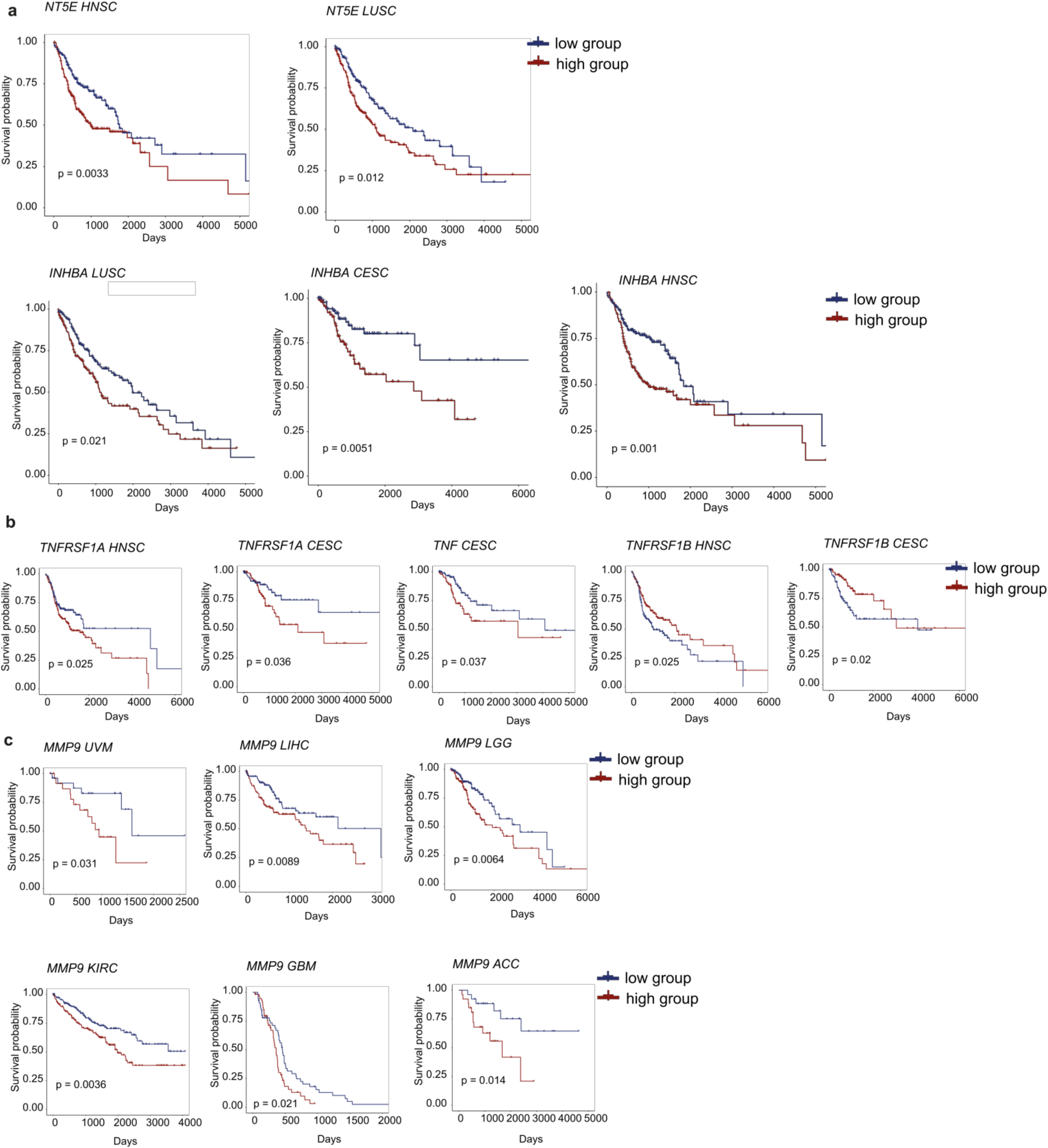
TSK mRNA levels correlate with overall survival in human SCC patients. a, TSK gene mRNA levels (*INHBA* and *NT5E*) correlate with shorter overall survival in human squamous cell carcinoma patients. LUSC, lung squamous cell carcinoma. HNSC, head and neck squamous cell carcinoma. CESC, cervical squamous cell carcinoma. All Kaplan-Meier plots show the overall survival, stratified by upper tertile (high) and bottom tertile (low) mRNA expression. P-value indicates a standard log-rank test. b, TNF-α, TNF-α receptor 1 and 2 mRNA level correlation with overall survival in human squamous cell carcinoma patients. c, *Mmp9* mRNA levels correlate with shorter overall survival in a total of 6 different cancer types. UVM, Uveal Melanoma. LIHC, Liver hepatocellular carcinoma. LGG, Brain Lower Grade Glioma. KIRC, Kidney renal clear cell carcinoma. GBM, Glioblastoma multiforme. ACC, Adrenocortical carcinoma.

**Extended Data Table 1. P4 and P60 sgRNA enrichments**.

**Extended Data Table 2. Pathway enrichment of the top 10 perturbations in P60 epidermal stem cells. Extended Data Table 3. WGCNA identifies 44 modules in the P60 skin**.

**Extended Data Table 4. TNF-**α **signaling is downregulated in human skin SCCs**.

**Extended Data Table 5. WGCNA identifies a TSK-like module within the set of 40 positively and negatively selected perturbations (Figure 5c)**.

